# Directed evolution of piperazic acid incorporation by a nonribosomal peptide synthetase

**DOI:** 10.1101/2023.04.03.535426

**Authors:** Philipp Stephan, Chloe Langley, Daniela Winkler, Jérôme Basquin, Lorenzo Caputi, Sarah E. O’Connor, Hajo Kries

## Abstract

Engineering of biosynthetic enzymes is increasingly employed to synthesize structural analogues of antibiotics. Of special interest are non-ribosomal peptide synthetases (NRPSs) responsible for production of important antimicrobial peptides. Here, directed evolution of an adenylation domain of a Pro-specific NRPS module completely switched substrate specificity to the non-standard amino acid piperazic acid (Piz) bearing a labile N-N bond. This success was achieved by LC-MS/MS based screening of small, rationally designed mutant libraries and can presumably be replicated with a larger number of substrates and NRPS modules. The evolved NRPS produces a Piz-derived gramicidin S analog. Thus, we give new impetus to the too-early dismissed idea that widely accessible low-throughput methods can switch the specificity of NRPSs in a biosynthetically useful fashion.

---

Non-ribosomal peptides (NRPs) and derivatives are an important class of antimicrobial molecules.^[1–3]^ The rising threat of antimicrobial resistance necessitates the generation and screening of novel bioactive compounds. NRPs are produced by enzyme complexes, so called non-ribosomal peptide synthetases (NRPS), working like assembly lines.^[2,4]^ On the assembly line, specialized domains organized in modules, one per amino acid, perform distinct biochemical activities.^[1]^ Adenylation domains (A-domains) recognize a specific amino acid, activate it under consumption of ATP, and load it onto the flexible phosphopantetheine arm of thiolation domains (T-domains). Condensation domains (C-domains) catalyze formation of a peptide bond between two adjacent T-domain-bound substrates. Thioesterase domains (TE-domains) release the peptide from the enzyme complex via hydrolysis or cyclisation. GrsA and GrsB are two NRPS proteins that together synthesize gramicidin S (GS, Figure 1a).^[5,6]^

**Figure 1.**
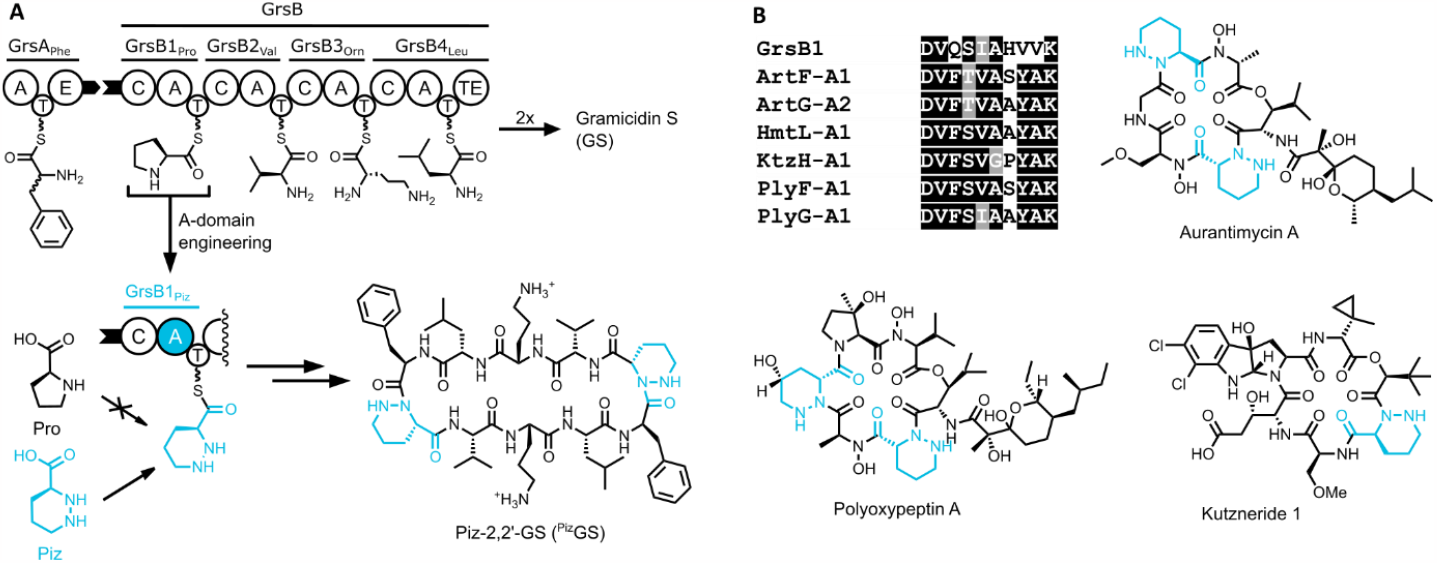
**A**. Biosynthesis of gramicidin S (GS) by the two NRPSs GrsA and GrsB.^[5,6]^ Through engineering of the A-domain of module GrsB1, _L_-Pro residues in GS are replaced with _L_-Piz, resulting in Piz-2,2’-GS (^Piz^GS). **B**. Alignment of specificity determining residues of GrsB1 with those of Piz-A-domains. The corresponding NRPs aurantimycin A^[35]^, polyoxypeptin A^[36]^, kutzneride 1^[37]^ contain Piz.

The modular architecture of NRPSs has inspired various engineering approaches to produce new or modified peptides.^[7–10]^ Often, attempts to replace entire modules, domains or smaller subdomains result in chimeric enzymes with reduced activity or premature termination of peptide synthesis.^[11–13]^ The discovery of a “specificity code” for A-domains^[14,15]^ was the basis for site-directed mutagenesis of the respective residues to change A-domain substrate specificity. Successful applications of this engineering strategy mainly enabled conservative substrate changes, or resulted in reduced enzyme activity^[16–19]^ with only a few examples of engineering allowing acceptance of non-natural amino acid substrates.^[20,21]^ Recently, directed evolution and high-throughput screening have been increasingly used in this context.^[22–27]^ Yeast surface display also has been employed to analyze large mutant libraries resulting in efficient recognition of β-amino acids and hydroxy acids. However, this technique requires a click-handle in the substrate and protein expression on the yeast surface, limiting wider applicability.^[26,27]^

Piz is an uncommon NRPS building block with only around 150 Piz-containing NRPs described (Figure 1a).^[28–30]^ In many of these, Piz is assumed to be responsible for bioactivity. Piz is a Pro analog with a six-membered ring incorporating a hydrazine, resulting in an even higher structural rigidity than Pro (Figure 1a).^[31]^ Piz is biosynthesized from ornithine and then incorporated into NRPs.^[32,33]^ The hydrazides of Piz provide a rare example of N-N bonds involved in peptide bond formation.^[34]^ Here, we have engineered the substrate specificity of the A-domain of the first module of the GS-producing NRPS GrsB (GrsB1) from Pro to Piz. After four rounds of evolution and screening of only 1,200 mutants for peptide production using LC-MS/MS, we generated a GrsB1 mutant showing a 140,000-fold improvement in Piz-preference.

To analyze Piz and Pro incorporation by module GrsB1 from the GS NRPS, we measured diketopiperazines (DKPs) that GrsB1 forms together with GrsA_Phe_ *in vitro* (Figure 2a). The A-domain of native module GrsB1_Pro_ shows minor DKP formation with Piz as substrate in addition to its main activity for Pro (Figure 2b). To guide our design efforts, we solved the structure of the apo-A_core_-domain of GrsB1 at a resolution of 2.6 Å (Figure 2c, Figure S1). Attempts to crystallize GrsB1 bound to either Piz or Pro were unsuccessful. We identified residues involved in substrate specificity of Piz-A-domains^[35–38]^ using NRPSpredictor2^[39]^ and compared these residues to the Pro-A-domain of GrsB1 (Figure 1b). Based on the differences in the active site, we created six mutants of GrsB1 to make Piz the preferred substrate. These proteins were Ni-affinity-purified from 400 mL of culture and tested *in vitro* for DKP production in the presence of equimolar Pro and Piz (10 mM). The product ratio was determined by UPLC-MS/MS and mutants conferring increased specificity for Piz were then also tested in combination (Figure S2). From these experiments inspired by natural Piz-A-domains, we thus obtained combination mutant GrsB1-AYA harboring three mutations (H755A, V763Y, V764A). While GrsB1-AYA shows 80-fold higher Piz specificity compared to GrsB1, Pro still remains the preferred substrate (Figure 2b and Figure S2) confirming once more that copying specificity codes alone is not sufficient to engineer A-domains.

**Figure 2.**
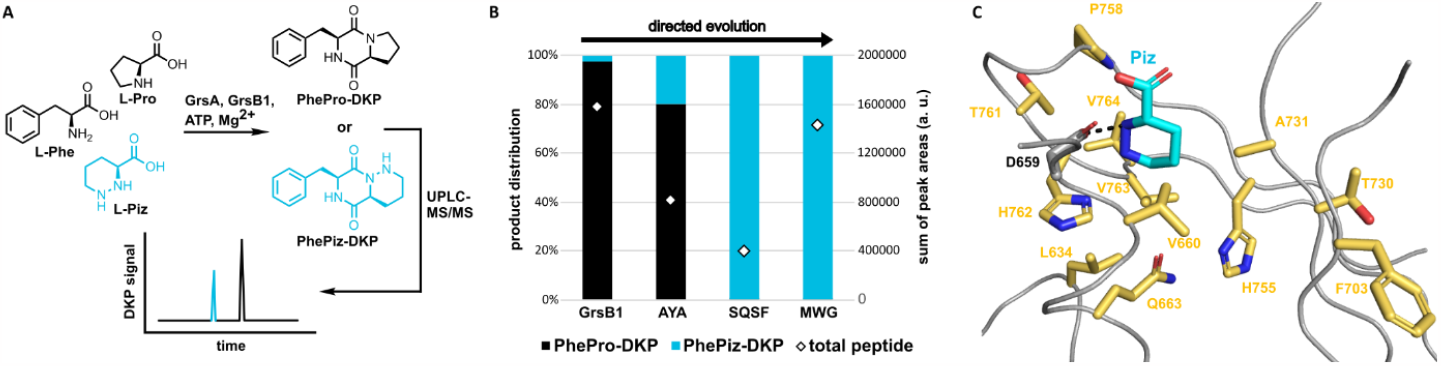
**A**. Diketopiperazine (DKP) formation with NRPS modules GrsA and GrsB1. When Pro and Piz are supplied in equimolar concentration as substrates for GrsB1, PhePro-DKP or PhePiz-DKP are formed. The ratio of the respective signals measured by UPLC-MS/MS can be used to compare Piz specificities of GrsB1 mutants. **B**. Directed evolution of Piz specificity. **C**. Structure of GrsB1 substrate binding pocket of GrsB1 solved to 2.6 Å resolution. To place the substrate in the active site, structure of GrsB1 was first aligned with the reported structure of PheA^[42]^ (PDB 1AMU) and visualized in Pymol. Next, the Phe ligand in 1AMU was replaced with Piz by aligning the α-amino and carboxyl groups. Residues that were mutated in this study (yellow) and the conserved Asp659 (grey) interacting with the α-amino group of Piz are shown as sticks.

To further improve Piz specificity, we considered positions within 5 Å of the substrate binding site that interact with the residues mutated in GrsB1-AYA (referred to as “second shell” residues). These positions revealed several less conserved differences between A-domains (Figure S3). To account for this ambiguity, we designed mutant libraries using partially degenerate codons covering all side-chain identities observed in the second shell of the Piz-A-domains. To screen these libraries, we performed the DKP assay with cell lysate of cultures grown in 96 well plates. Amino acids, ATP, and Ni-affinity-purified GrsA were supplied to lysate containing GrsB1 mutants. By UPLC-MS/MS, peptides from hundreds of variants could be quantified in parallel. By screening 264 mutants, we identified GrsB1-SQSF, harboring four mutations in addition to those of GrsB1-AYA (T730S, P758Q, T761S, H762F), and showing perfect specificity for Piz (Figure 2b). However, the overall production of DKP was reduced (Figure S4).

To improve the yield of Piz-derived peptide, we performed saturation mutagenesis with NNK codons^[40]^ on 13 residues neighboring previously mutated positions and combined the hits in another round of screening. Screening in lysate for DKP identified variant GrsB1-MWG carrying five additional mutations (L634V, V660M, Q663W, F703M, A731G [Figure S5, Figure S6]). GrsB1-MWG not only has perfect Piz specificity like GrsB1-SQSF, but also DKP formation activity comparable to GrsB1 with Pro (Figure 2b). While GrsB1 incorporates 0.5% Piz with equimolar Piz and Pro offered as substrates, GrsB1-MWG has no detectable Pro activity demonstrating excellent molecular recognition by the engineered enzyme. To better understand the importance of the mutations, we attempted to crystallize both apo- and substrate-bound forms of the GrsB1-MWG mutant but were unsuccessful. However, mapping the mutations on the structure of GrsB1 revealed residues controlling Piz substrate specificity to be distributed throughout the binding pocket, suggesting extensive remodeling of the active site (Figure 2c). Homology modelling of GrsB1-MWG based on the GrsB1 crystal structure and docking of Piz identified the potential role of mutation V763Y to hydrogen-bond with the distal nitrogen atom of Piz, aiding substrate orientation (Figure S7b). In addition, mutation P758Q may increase flexibility of the neighboring β13β14 loop, the conformation of which is important for β-amino acid recognition.^[26,41]^

We determined the thermal stability of representative variants by thermal shift assays (Figure S8). GrsB1-SQSF is the least thermally stable enzyme with a melting point (T_m_) of 40.7 °C, lowered from 46.4 °C for GrsB1, perhaps explaining its lower activity. The additional mutations in GrsB1-MWG mostly recovered protein stability (T_m_ = 43.8 °C). These findings correlate with the observed protein yield after Ni-affinity purification. Yields for GrsB1 and GrsB1-MWG are comparable (volumetric yield: 51 – 62 mg/L) whereas GrsB1-SQSF shows lower production of soluble enzyme (38 mg/L, Figure S9).

To compare the affinity of GrsB1 and GrsB1-MWG for their ligands, we titrated the proteins with stable adenosyl monosulfamate (AMS) analogues^[43,44]^ of the activated adenylates Pro-AMP and Piz-AMP and measured T_m_ values. The change in T_m_ was plotted against the ligand concentration to determine dissociation constants (*K*_D_’s, Figure S10-12). Since Piz-AMS synthesis proved difficult, nipecotic acid-AMS (Nip-AMS) was used instead. Piz and the β-amino acid Nip both have a six-membered ring as a side-chain, but in Nip, the nitrogen atom in the α-position of Piz is replaced with a carbon atom. The *K*_D_ for Nip-AMS increases slightly from GrsB1 (14 μM) to GrsB1-MWG (52 μM), while the *K*_D_ for Pro-AMS increases dramatically by 2000-fold (0.7 μM and 1.5 mM, respectively). Hence, a switch from 20-fold Pro-AMS to 29-fold Nip-AMS preference has been achieved by evolving for Piz preference. The connection between Piz and Nip specificity probably indicates that the β-amino group of Nip and the corresponding nitrogen atom in Piz play an analogous and important role in molecular recognition.

In addition, we measured adenylation kinetics for Pro and Piz with the GrsB1 variants obtained during directed evolution. Adenylation kinetics were recorded using the MesG assay that enzymatically couples the rate of pyrophosphate (PP_i_) release during adenylation to the production rate of a photometrically detectable nucleoside analogue (Table 1 and Figure S13).^[45]^ Addition of hydroxylamine prevents accumulation of the inhibitory amino acyl-adenylates.^[45,46]^ As expected, GrsB1 and GrsB1-MWG have the highest catalytic efficiency (*k*_cat_/*K*_M_) for the native substrate Pro (17.4 min^-1^ mM^-1^) and the target substrate Piz (1.2 min^-1^ mM^-1^), respectively. Interestingly, all mutants maintain similar or even higher turnover rates (*k*_cat_) for Piz between 2.8 and 5.0 min^-1^ that are all well in the range of typical peptide formation rates of NRPSs.^[47,48]^ However, the Michaelis-constant (*K*_M_) dramatically increases for Pro from 0.14 mM (GrsB1) to 10 mM (GrsB1-AYA) after the first round of directed evolution, and then becomes undetectable in subsequent rounds. In contrast, the *K*_M_ for Piz drops from 9.2 mM in GrsB1 to 2.4 mM in GrsB1-MWG. Altogether, directed evolution has had a dramatic impact on the molecular recognition of the structurally similar substrates Pro and Piz while maintaining biosynthetically useful catalytic rates.

**Table 1.** Adenylation kinetics of representative GrsB1 mutants.*

| | | $K_M$<br>(mM) | $k_{cat}$<br>(min<sup>-1</sup>) | $k_{cat}/K_M$<br>(mM<sup>-1</sup> min<sup>-1</sup>) |
| --- | --- | --- | --- | --- |
| **GrsB1-WT** | L-Pro | 0.14 ± 0.03 | 2.4 ± 0.3 | 17.4 |
|  | L-Piz | 9.2 ± 0.9 | 4.1 ± 0.2 | 0.4 |
| **GrsB1-AYA** | L-Pro | 10 ± 2 | 1.8 ± 0.2 | 0.2 |
|  | L-Piz | 17 ± 4 | 5.0 ± 0.9 | 0.3 |
| **GrsB1-SQSF** | L-Pro | n.d.<sup>#</sup> | n.d. | n.d. |
|  | L-Piz | 4.4 ± 0.6 | 2.9 ± 0.2 | 0.7 |
| **GrsB1-MWG** | L-Pro | n.d. | n.d. | n.d. |
|  | L-Piz | 2.4 ± 0.3 | 2.8 ± 0.1 | 1.2 |

\*Adenylation kinetics were determined using the MesG/hydroxylamine assay. Initial velocities were plotted against substrate concentrations according to the Michaelis-Menten model to obtain values for $K_M$ and $k_{cat}$ . Each enzyme was measured as biological duplicate. The errors indicate the error of fit. <sup>#</sup>n.d.: not detectable.

After successfully switching the specificity of GrsB1, we used the engineered NRPS module to produce ^Piz^GS in *E. coli*. For this purpose, the *grsB1* gene in *grsB* was replaced with the engineered *grsB1-MWG* in plasmid pSU18-grsTAB carrying the entire GS biosynthetic gene cluster.^[49]^ The gene cluster was then expressed heterologously in *E. coli* following a previously established protocol^[49]^ while supplementing Piz to the media. Surprisingly, the evolutionary intermediate SQSF-VM showed the highest production yield for ^Piz^GS under *in vivo* conditions, not the mutant GrsB1-MWG. In a brief optimization of production conditions (Figure S14), we found that media composition, Piz concentration, shaking speed, and the ratio of culture to flask volume strongly influence the production rate. Using 50 mL LB media in 2 L culture flasks shaking at 400 rpm, we obtained 15 mg/L of ^Piz^GS in extracts of the cell pellet while no production of native GS was detectable. Concentrations were estimated by quantification of UPLC-MS/MS signals using a GS standard assuming that GS and ^Piz^GS ionize with similar efficiency. ^Piz^GS was identified by detecting a specific mass transition 586>86 in MRM mode ([M+2H]^2+^ for ^Piz^GS parent ion to iminium ion of Piz side chain after decarboxylation as daughter ion, Figure 3a) as well as measuring HRMS (calculated: *m/z* = 586.3712 [M+2H]^2+^ for C_60_H_96_N_14_O_10_^2+^, measured: *m/z* = 586.3713 [M+2H]^2+^, Figure 3c). Compared to native GS, ^Piz^GS production reached a level of 20% (Figure 3b). When no exogenous Piz is added to the production culture, neither ^Piz^GS nor native GS are produced, highlighting again the high selectivity for Piz over Pro in the evolved GrsB1.

**Figure 3.**
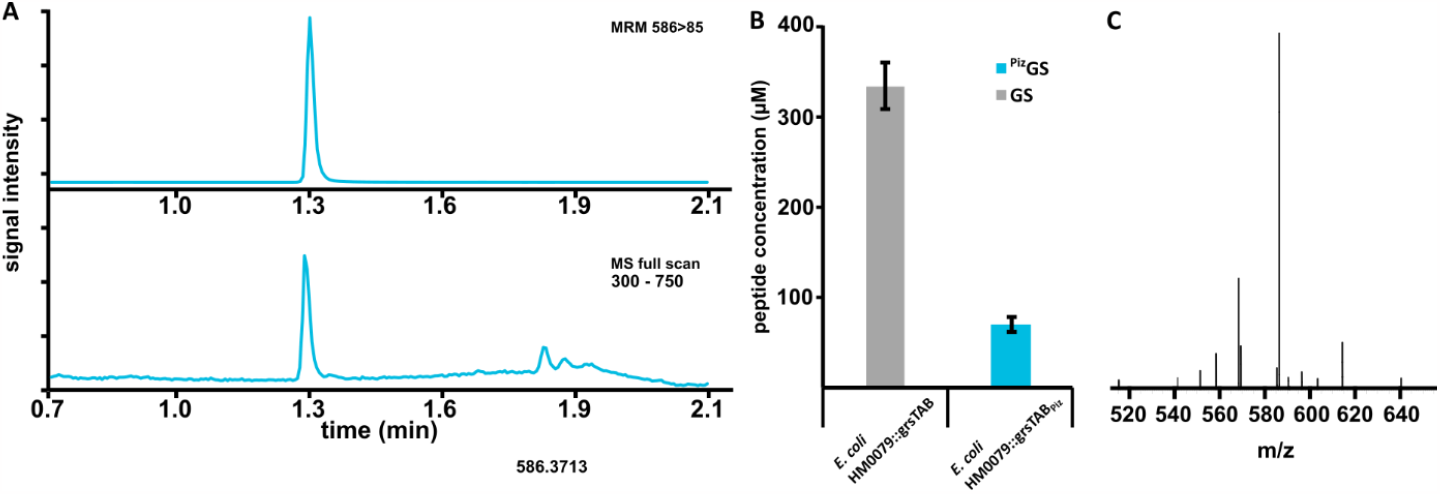
**A**. Total ion chromatograms for full MS scan and ^Piz^GS specific MRM (586>85) for crude extracts of cell pellets from ^Piz^GS producing *E. coli* cultures. Both traces are normalized to the highest peak. Using MRM measurement specific for native GS no signals for the compound could be detected. **B**. Comparison of product yield in pellet extracts for cultures producing native GS (*E. coli* HM0079::grsTAB) or ^Piz^GS (*E. coli* HM0079::grsTAB_Piz_) grown under optimized conditions (TB medium, 6 mM ornithine, 30 °C, 5 days or LB medium, 6 mM ornithine, 15 mM Piz, 30 °C, 5 days, respectively). A GS standard of known concentration was used to calculate concentrations. ^Piz^GS concentration was estimated assuming that GS and ^Piz^GS ionize comparably well. Error bars indicate standard deviation from three biological replicates. **C**. HRMS spectra for purified ^Piz^GS (calculated: *m/z* = 586.3712 [M+2H]^2+^, measured: *m/z* = 586.3713 [M+2H]^2+^, Δ = 0.2 ppm).

Directed evolution is a powerful tool to engineer tailor-made enzyme catalysts.^[50–52]^ Here, we have applied a directed evolution strategy on an A-domain of the GS biosynthetic assembly line to incorporate the non-proteinogenic amino acid Piz. Inspired by natural specificity-codes we performed rational mutagenesis and further refined the resulting variants using random mutagenesis. Thus, we achieved a complete switch in substrate specificity after screening only 1200 mutants in total. Interestingly, GrsB1-SQSF, the first mutant showing a complete switch in substrate specificity (Figure 2b), carries mutations in positions previously shown to induce selectivity for β-amino acids or α-hydroxy acids (GrsB1: P758Q, T761S, H762F; TycA_βPhe_: T761C, I762L; TycA_pPLA_: P758L; numbering of TycA_βPhe_ and TycA_pPLA_ respective to GrsB1).^[26,27]^ Considering that the main distinction between Pro and Piz is the additional nitrogen in a “β-like’’ position of Piz, it might be expected that positions influencing β/α specificity could also impact Piz/Pro specificity. Replacing Pro’s amino group with a hydrazine in Piz may have been expected to affect the condensation reaction as strongly as the adenylation reaction. Since the “α-like” nitrogen of Piz used for peptide bond formation is only weakly nucleophilic,^[53,54]^ it was speculated that C-domains might have to co-evolve with A-domains to achieve Piz incorporation into peptides.^[30]^ However, efficient Piz incorporation into GS suggests that the unchanged C-domain does not discriminate against Piz. Therefore, replacing Pro with Piz in other NRPs may be feasible. Piz incorporation may enable further modification of natural products if the N-N-bond in Piz can be chemically cleaved, which would convert Piz into Orn.

Previous NRPS engineering attempts have failed to achieve consistent results based on A-domain specificity codes alone.^[16–19]^ While transplanting the specificity code did not yield a satisfactory result here, when combined with random mutagenesis, we were able to selectively switch substrate specificity. The resulting specificity determining code (DMWSIGAYAK) is close but not identical to the consensus of natural codes identified by Wei *et al*.,^[30]^ for Piz-A-domains (DVFSVAxYAK). The second and third position are highly conserved in natural specificity codes (Val and Phe) but different here (Met and Trp).

These differences are likely due to context dependence of the specificity code and highlight the need for unbiased mutant screening for successful directed evolution of A-domains.

Yeast-surface display has been previously used to solve an A-domain engineering task of similar difficulty, the conversion of a Phe-A-domain into a βPhe-A-domain.^[26]^ However, these efforts require a specialized high throughput assay that is somewhat limited in scope and time-consuming to implement. In this study, we have directly measured peptide formation to prevent optimizing partial reactions at the expense of the overall functionality.^[25]^ Screening assays capable of analyzing mutants in 96-well plate format under native conditions, such as the LC-MS/MS assay used here, could be implemented for many other NRPSs.

The rise in antibiotic resistant bacteria makes the discovery of new or modified compounds a primary concern. Our highly efficient and broadly applicable screen can help in this regard by enabling faster and easier engineering strategies for NRPSs, an important source for bioactive peptides.

## Acknowledgements

We would like to thank Farzaneh Pourmasoumi, Oliver Waldmann, and Katharina Heise for helpful discussions and technical assistance. We acknowledge financial support by the Daimler und Benz Stiftung, the Fonds der Chemischen Industrie, and the International Leibniz Research School (ILRS).

## References

[1] S. A. Sieber, M. A. Marahiel, Chem. Rev. 2005, 105, 715–738.

[2] R. D. Süssmuth, A. Mainz, Angew. Chem. Int. Ed. 2017, 56, 3770–3821.

[3] Y. Liu, S. Ding, J. Shen, K. Zhu, Nat. Prod. Rep. 2019, 36, 573–592.

[4] J. Grünewald, M. A. Marahiel, Microbiol Mol Biol Rev 2006, 70, 121–146.

[5] J. Krätzschmar, M. Krause, M. A. Marahiel, J Bacteriol 1989, 171, 5422–5429.

[6] F. Saito, K. Hori, M. Kanda, T. Kurotsu, Y. Saito, The Journal of Biochemistry 1994, 116, 357–367.

[7] R. H. Baltz, ACS Synth. Biol. 2014, 3, 748–758.

[8] E. Kim, B. S. Moore, Y. J. Yoon, Nat Chem Biol 2015, 11, 649–659.

[9] A. S. Brown, M. J. Calcott, J. G. Owen, D. F. Ackerley, Nat. Prod. Rep. 2018, 35, 1210–1228.

[10] K. A. J. Bozhüyük, A. Linck, A. Tietze, J. Kranz, F. Wesche, S. Nowak, F. Fleischhacker, Y.-N. Shi, P. Grün, H. B. Bode, Nat. Chem. 2019, 11, 653–661.

[11] T. Stachelhaus, A. Schneider, M. A. Marahiel, Science 1995, 269, 69–72.

[12] M. J. Calcott, J. G. Owen, I. L. Lamont, D. F. Ackerley, Appl Environ Microbiol 2014, 80, 5723–5731.

[13] H. Kries, D. L. Niquille, D. Hilvert, Chemistry & Biology 2015, 22, 640–648.

[14] T. Stachelhaus, D. Mootz, A. Marahiel, Chemistry & Biology 1999, 6, 493–505.

[15] G. L. Challis, J. Ravel, C. A. Townsend, Chemistry & biology 2000, 7, 211–224.

[16] K. Eppelmann, T. Stachelhaus, M. A. Marahiel, Biochemistry 2002, 41, 9718–9726.

[17] C.-Y. Chen, I. Georgiev, A. C. Anderson, B. R. Donald, Proc. Natl. Acad. Sci. U.S.A. 2009, 106, 3764–3769.

[18] J. Thirlway, R. Lewis, L. Nunns, M. Al Nakeeb, M. Styles, A. W. Struck, C. P. Smith, J. Micklefield, Angew. Chem. Int. Ed. 2012, 51, 7181–7184.

[19] J. W. Han, E. Y. Kim, J. M. Lee, Y. S. Kim, E. Bang, B. S. Kim, Biotechnol Lett 2012, 34, 1327–1334.

[20] H. Kries, R. Wachtel, A. Pabst, B. Wanner, D. Niquille, D. Hilvert, Angew. Chem. Int. Ed. 2014, 53, 10105–8.

[21] H. Kaljunen, S. H. H. Schiefelbein, D. Stummer, S. Kozak, R. Meijers, G. Christiansen, A. Rentmeister, Angew. Chem. Int. Ed. 2015, 54, 8833–8836.

[22] M. A. Fischbach, J. R. Lai, E. D. Roche, C. T. Walsh, D. R. Liu, Proc. Natl. Acad. Sci. U.S.A. 2007, 104, 11951–11956.

[23] B. Villiers, F. Hollfelder, Chemistry & Biology 2011, 18, 1290–1299.

[24] B. S. Evans, Y. Chen, W. W. Metcalf, H. Zhao, N. L. Kelleher, Chemistry & Biology 2011, 18, 601–607.

[25] K. Zhang, K. M. Nelson, K. Bhuripanyo, K. D. Grimes, B. Zhao, C. C. Aldrich, J. Yin, Chemistry & Biology 2013, 20, 92–101.

[26] D. L. Niquille, D. A. Hansen, T. Mori, D. Fercher, H. Kries, D. Hilvert, Nature Chem 2018, 10, 282–287.

[27] A. Camus, G. Truong, P. R. E. Mittl, G. Markert, D. Hilvert, J. Am. Chem. Soc. 2022, 144, 17567–17575.

[28] J. Oelke, D. J. France, T. Hofmann, G. Wuitschik, S. V. Ley, Natural Product Reports 2011, 28, 1445–1471.

[29] K. D. Morgan, R. J. Andersen, K. S. Ryan, Nat. Prod. Rep. 2019, 36, 1628–1653.

[30] Z.-W. Wei, H. Niikura, K. D. Morgan, C. M. Vacariu, R. J. Andersen, K. S. Ryan, J. Am. Chem. Soc. 2022, 144, 13556–13564.

[31] N. Xi, L. B. Alemany, M. A. Ciufolini, J. Am. Chem. Soc. 1998, 120, 80–86.

[32] S. Neumann, W. Jiang, J. R. Heemstra, E. A. Gontang, R. Kolter, C. T. Walsh, ChemBioChem 2012, 13, 972–976.

[33] Y. L. Du, H. Y. He, M. A. Higgins, K. S. Ryan, Nature Chemical Biology 2017, 13, 836–838.

[34] A. J. Waldman, T. L. Ng, P. Wang, E. P. Balskus, Chem. Rev. 2017, 117, 5784–5863.

[35] H. Zhao, L. Wang, D. Wan, J. Qi, R. Gong, Z. Deng, W. Chen, Microb Cell Fact 2016, 15, 160.

[36] Y. Du, Y. Wang, T. Huang, M. Tao, Z. Deng, S. Lin, BMC Microbiology 2014, 14, 1–12.

[37] G. Fujimori, S. Hrvatin, C. S. Neumann, M. Strieker, M. A. Marahiel, C. T. Walsh, Proc. Natl. Acad. Sci. U.S.A. 2007, 104, 16498–16503.

[38] J. Ma, Z. Wang, H. Huang, M. Luo, D. Zuo, B. Wang, A. Sun, Y.-Q. Cheng, C. Zhang, J. Ju, Angew. Chem. Int.Ed. 2011, 50, 7797–7802.

[39] M. Röttig, M. H. Medema, K. Blin, T. Weber, C. Rausch, O. Kohlbacher, Nucleic Acids Research 2011, 39, 362–367.

[40] K. Miyazaki, F. H. Arnold, J Mol Evol 1999, 49, 716–720.

[41] A. Miyanaga, J. Cieślak, Y. Shinohara, F. Kudo, T. Eguchi, Journal of Biological Chemistry 2014, 289, 31448–31457.

[42] Conti, T. Stachelhaus, M. a Marahiel, P. Brick, Embo J 1997, 16, 4174–4183.

[43] H. Ueda, Y. Shoku, N. Hayashi, J. Mitsunaga, Y. In, M. Doi, M. Inoue, T. Ishida, Biochimica et Biophysica Acta (BBA) - Protein Structure and Molecular Enzymology 1991, 1080, 126–134.

[44] R. Finking, A. Neumüller, J. Solsbacher, D. Konz, G. Kretzschmar, M. Schweitzer, T. Krumm, M. A. Marahiel, ChemBioChem 2003, 4, 903– 906.

[45] D. J. Wilson, C. C. Aldrich, Analytical Biochemistry 2010, 404, 56–63.

[46] E. J. Drake, B. R. Miller, C. Shi, J. T. Tarrasch, J. A. Sundlov, C. Leigh Allen, G. Skiniotis, C. C. Aldrich, A. M. Gulick, Nature 2016, 529, 235–238.

[47] A. Stanišić, A. Hüsken, P. Stephan, D. L. Niquille, J. Reinstein, H. Kries, ACS Catal. 2021, 11, 8692–8700.

[48] I. Folger, N. Frota, A. Pistofidis, D. Niquille, D. Hansen, T. M. Schmeing, D. Hilvert, High-Throughput Reprogramming of an NRPS Condensation Domain, In Review, 2023.

[49] F. Pourmasoumi, S. Hengoju, K. Beck, P. Stephan, L. Klopfleisch, M. Hoernke, M. A. Rosenbaum, H. Kries, Screening Megasynthetase Mutants at High Throughput Using Droplet Microfluidics, Biorxiv, 2023.

[50] M. S. Packer, D. R. Liu, Nat Rev Genet 2015, 16, 379–394.

[51] F. H. Arnold, Angew. Chem. Int. Ed. 2018, 57, 4143–4148.

[52] Y. Wang, P. Xue, M. Cao, T. Yu, S. T. Lane, H. Zhao, Chem. Rev. 2021, 121, 12384–12444.

[53] M. A. Ciufolini, N. Xi, Chem. Soc. Rev. 1998, 27, 437.

[54] M. A. Ciufolini, T. Shimizu, S. Swaminathan, N. Xi, Tetrahedron Letters 1997, 38, 4947–4950.

