## Supporting Information for "Directed evolution of piperazic acid incorporation by a nonribosomal peptide synthetase"

[c] Dr. J. Basquin
Department of Structural Cell Biology
Max Planck Institute for Biochemistry
Am Klopferspitz 18, Martinsried 82152 Planegg, Germany

### 1. Cloning

#### 1.1. General cloning

Cloning was carried out in *E. coli* strain Stellar (Takara Bio Europe). Holo proteins were expressed in *E. coli* strain HM0079[1]. For the purification of plasmid DNA, DNA fragments, and PCR products, NucleoSpin Plasmid, and Gel and PCR clean-up kits (Macherey Nagel) were used. DNA fragments were amplified with Q5 polymerase (New England Biolabs, Massachusetts) following the supplier's instructions. Assembly of PCR fragments containing vector-specific overhangs and linearized vector was done using the InFusion cloning kit (Takara Bio Europe). Oligonucleotides used as primers (Table S1) were made by custom synthesis (Eurofins Genomics) and sequence confirmation of assembled constructs was done by Sanger sequencing (GENEWIZ).

#### 1.2. Plasmids

##### 1.2.1. pTrc99a-GrsB1_KpnI

To make plasmid pTrc99a-GrsB1_KpnI, an additional KpnI cut site was introduced into plasmid pTrc99a-GrsB1_corr_CAT[2] by creating a silent mutation at the codon translated as GrsB1-V969. Plasmid pTrc99a-GrsB1_corr_CAT was linearized with AflII and BamHI and the mutated GrsB1 assembled in two overlapping fragments that were amplified by PCR using primer pairs pTrc99a-GrsB1_V868KpnI_fw / pTrc99a-GrsB1_BamHI_rev and GrsB1_piz_AflII_fw / pTrc99a-GrsB1_V969_rev.

##### 1.2.2. pOPINF-GrsB1-ANTsub and pOPINF-MWG-ANTsub

For crystallization trials, only the large N-terminal $A_{core}$-domains of GrsB1 or GrsB1-MWG were expressed[3]. For this purpose, the corresponding genes were cloned into vector pOPINF (OPPF-UK, Addgene #26042) linearized with KpnI and HindIII. The corresponding regions on genes *grsB1* or *grsB1-MWG* were amplified by PCR using primer pair pOPINF-GrsB1-A_3_fw / pOPINF_GrsB1-A-NTsub_rev.

##### 1.2.3. GrsB1 active site mutants

Plasmids encoding active site mutants of GrsB1 were cloned by assembling two PCR amplified DNA fragments of the *grsB1_KpnI* gene with compatible overhangs in vector pTrc99a-GrsB1_KpnI linearized with AflII and KpnI, which are located upstream and downstream of the mutated fragment, respectively. Fragment 1 consisted of the region between the AflII site and the position to be mutated. Fragment 2 consisted of the region between the site of mutation and the KpnI site. Forward primers for fragment 2 always carried the mutations to be introduced.

##### 1.2.4. GrsB1 second shell libraries

Libraries of GrsB1 were cloned using primers with degenerate codons (Metabion) to either include a specific subset of mutations or NNK codons for full saturation mutagenesis. GrsB1 libraries were amplified by PCR in two fragments similar to the active site mutants with overlapping ends and assembled by InFusion cloning into plasmid pTrc99a-GrsB1_KpnI linearized with AflII and KpnI.

##### 1.2.5. pSU18-grsTAB$_{Piz}$

For production of $^{Piz}$GS, the gene of a GrsB1 mutant was cloned into plasmid pSU18-grsTAB following a previously established cloning strategy for the GS cluster[4]. Plasmids pTrc99a-GrsB1-SQSF-VM and plasmids pTrc99a-GrsB1-MWG were used as template for PCR amplification with primer pair GrsB1_piz_AflII_fw / GrsB1_piz_KpnI_2_rev. The fragment was cloned into plasmids pTrc99a-grsAB1[4] linearized with AflII and KpnI. The resulting intermediary plasmids were used as template for PCR amplification using primer pair GrsAB1_grsTAB_fw / GrsAB1_grsTAB_rev and the PCR products cloned into plasmid pSU18-grsTAB linearized with EcoNI to result in pSU18-grsTAB$_{Piz}$ and pSU18-grsTAB$_{MWG}$, respectively.

### 2. Protein production and purification

#### 2.1. General protein production and purification protocol

In 2 L Erlenmeyer flasks, 400 ml of 2xYT liquid media containing the appropriate antibiotic were inoculated with 400 µl of a densely grown preculture of *E. coli* HM0079 carrying the respective plasmid with the gene of the protein to be produced. The culture was incubated at 37 °C with agitation at 250 rpm until the culture reached an $OD_{600}$ of 0.8 after ca. 4 h. The shaker was cooled to 18 °C and the culture incubated for 30 min. Protein production was induced with 375 µM IPTG (150 µl of a 1 M stock solution) and the culture incubated at 18 °C for 18 h with continuous shaking at 250 rpm. Cells were harvested by centrifugation at 8,000 g for 5 min and resuspended in 15 ml buffer A1 (50 mM Tris pH 7.4, 500 mM NaCl, 20 mM imidazole, 2 mM TCEP). 100 µl protease inhibitor cocktail (Sigma, P8849) was added and cells lysed by sonication. The lysate was cleared by centrifugation at 50,000 g for 30 min at 6 °C. The supernatant was applied to 2 ml Ni-IDA suspension (Rotigarose, Roth) equilibrated with buffer A1. The column was washed with buffer A1 (3 x 5 ml) to remove unspecifically bound proteins. The protein of interest was eluted in 1 ml fractions with buffer B1 (50 mM Tris pH 7.4, 500 mM NaCl, 300 mM imidazole, 2 mM TCEP). Samples of each fraction were taken for SDS-PAGE analysis and the concentration determined by measuring absorption at 280 nM using a Take3 plate on an Epoch2 microplate reader using calculated extinction coefficients. Protein containing fractions were pooled, buffer exchanged to storage buffer (100 mM HEPES pH 8, 150 mM NaCl, 5% [v/v] glycerol) and adjusted to a final concentration of 50 µM using Vivaspin centrifugal filters with 50 kDa MWCO (Sartorius). Protein stocks were flash frozen in liquid nitrogen and stored at -80 °C.

#### 2.2. Production of 3C protease

3C protease was produced from *E. coli* BL21(DE3)::pE28a-HRV-3C mainly following the general protocol for protein purification with the only difference that no protease inhibitor cocktail was added to the resuspended cells before cell lysis. After Ni-NTA purification, 3C protease was adjusted to 5 mg/ml in storage buffer, aliquoted, and kept at -80 °C.

#### 2.3. Protein production for crystallography

Production of $A_{core}$ domains for crystallization was carried out using *E. coli* KRX (Promega) cells. 400 ml of 2xYT media with the appropriate antibiotic were inoculated with 1 ml of a densely grown overnight culture. Cells were grown at 37 °C, 250 rpm for 4 h until the culture reached an $OD_{600}$ of 0.8. Temperature was reduced to 18 °C and the culture incubated for 30 min before being induced with 2 ml 20% (w/v) rhamnose and 400 µl 1 M IPTG. Protein production was carried out for 18 h at 18 °C, shaking at 250 rpm. Ni-NTA purification followed the general protocol. Afterwards, the protein was digested with 3C protease to remove the N-terminal $His_6$-tag. The protease was added at a 1:50 (w/w) ratio to the protein and the digest incubated at 6 °C for 16 h. The cleaved $His_6$-tag as well as 3C protease were removed from the protein of interest by applying the digest to a Bio-Scale Mini Nuvia IMAC 5 ml cartridge (Bio-Rad) connected to a Bio-Rad NGC Chromatography system and eluting with a gradient of increasing imidazole concentration. Protein containing fractions were pooled and buffer exchanged to low-salt buffer (20 mM Tris pH 8, 20 mM NaCl) and applied to anion exchange chromatography using a Capto HiRes Q 5/50 column (Cytiva) eluting with a gradient of NaCl (increasing from 20 to 300 mM). Again, protein containing fractions were pooled, buffer exchanged to storage buffer (100 mM HEPES pH 8, 150 mM NaCl, 5% (v/v) glycerol) and adjusted to a final concentration of 20 mg/ml.

#### 2.4. SDS-PAGE of purified proteins

Purity of proteins was determined by SDS-PAGE (Figure S9) using Bolt 4-12% Bis-Tris Plus Gels (ThermoFisher Scientific) with MES-SDS running buffer (Novex). Triple Color Protein Standard III (Serva) was run alongside the protein samples as a size standard. The gels were run at 200 V for 22 min and stained with Quick Coomassie stain (Serva).

### 3. *In vitro* peptide formation assays

#### 3.1. Diketopiperazine formation assay

To make diketopiperazines (DKPs), 5 µM of GrsB1 or the respective GrsB1 mutants were mixed with 1 µM GrsA in a reaction buffer containing 20 mM HEPES, 0.5 mM $MgCl_2$, 1 µM TCEP, 1 mM L-Phe (Roth), 10 mM L-Pro (Roth) and 10 mM L-Piz. Reactions were started by addition of 5 mM ATP and incubated at 37 °C for 1 h. Afterwards, they were quenched by heat through incubation at 95 °C for 5 min. Denatured proteins were removed by centrifugation at 20,000 g for 5 min. The resulting supernatant was diluted 1:50 in 15% (v/v) MeCN for UPLC-MS/MS analysis.

##### 3.1.1. UPLC-MS/MS conditions

Chromatography was performed on a Waters ACQUITY H-class UPLC system (Waters) with an injection volume of 2 µL. Acetonitrile (A) and water with 0.1 % formic acid (B) were used as strong and weak eluent, respectively. Separation of products was achieved on a Cortecs UPLC C18 column (1.6 µm, 2.1 x 50 mm) with a linear gradient of 2-50 % A over 1.2 min (flow rate 0.5 mL/min) followed by 0.5 min wash with 100% A and 1.1 min re-equilibration. Acetonitrile was used as a needle wash solvent between the samples. Data acquisition and quantitation were done using the MassLynx and TargetLynx software (version 4.1).

MS/MS analyses were performed on a Xevo TQ-S micro (Waters) tandem quadrupole instrument with ESI ionisation source in positive ion mode. Nitrogen was used as desolvation gas and argon as collision gas. The following source parameters were used: capillary voltage 0.5 kV, cone voltage 4 V, desolvation temperature 600 °C, desolvation gas flow 1000 L/h. PhePro-DKP and PhePiz-DKP were detected via the 245.29>119.99 and 260.33>120.04 transitions, respectively, recorded in multiple reaction monitoring (MRM) mode.

#### 3.2. 96 well lysate screening assay

Plasmids encoding for GrsB1 libraries were transformed into *E. coli* HM0079 by heat shock. The transformation mixture was diluted and plated onto LB + amp plates to obtain about 100 cfu per plate. Sterile 96-well plates (Sarstedt) were prepared with 150 µl 2xYT media containing the appropriate antibiotic per well, and each well was inoculated with one colony. Colonies of *E. coli* HM0079 producing either GrsB1, GrsB1-AYA, or GrsA were used as controls distributed over the plate. 96-well plates were incubated for 18 h at 37 °C and 400 rpm to be used as pre-culture.

MegaBlock 2.2 ml 96-deepwell plates (Sarstedt) were prepared with 1 ml 2xYT media per well containing the appropriate antibiotic. Each well was inoculated with 70 µl from the preculture plate. Cultures were grown for 4 h at 37 °C and 400 rpm. The temperature was then lowered to 18 °C and the plates incubated for 30 min. Protein production was induced by addition of 2 mM IPTG in 2xYT media to a final concentration of 0.25 mM IPTG per well. Protein production was carried out at 18 °C and 400 rpm for 18 h. Cells were harvested by centrifugation at 4,000 g and 6 °C for 30 min. The supernatant was removed, the pellet resuspended in lysis buffer (20 mM HEPES pH 8, 50 mM NaCl, 1 mM EDTA, 1.5 mg/ml lysozyme), and incubated at room temperature for 30 min. Cells were completely lysed by slowly freezing the cell suspension overnight at -20 °C and thawing it the next day at room temperature for 3 h. To the cell lysate, DNA removal mix (20 mM HEPES pH 8, 50 mM NaCl, 1 mM EDTA, 2 mM $MgCl_2$, 2 mM TCEP, 3 U/ml Turbonuclease [Jena Bioscience]) was added and the lysate incubated for 20 min on ice. Cell debris was removed by centrifugation at 4,000 g for 30 min at 6 °C. The supernatant was diluted 1:10 with *dd*$H_2O$ and then mixed 1:1 with 2 x DKP assay buffer (40 mM HEPES pH 8, 1 mM $MgCl_2$, 2 mM TCEP, 10 mM ATP, 2 mM $d_5$-L-Phe [CDN Isotopes], 20 mM L-Pro, and 20 mM L-Piz, 2 µM GrsA) in a 96 well PCR plate and incubated for 1 h at 37 °C in a thermocycler and quenched afterwards at 95 °C for 5 min. Denatured proteins were removed by centrifugation. The supernatant was diluted 1:20 with 15% (v/v) MeCN + 0.1% (v/v) formic acid in a 384 well plate (Brandt) covered with aluminium foil and analysed by UPLC-MS/MS. Samples were separated on a Cortecs UPLC C18 column (1.6 µm, 2.1 x 30 mm) with a linear gradient of 10-15% A over 1.2 min (flow rate 0.5 mL/min) followed by 0.5 min wash with 100 % A and 0.5 min re-equilibration. $d_5$-PhePro-DKP and $d_5$-PhePiz-DKP were detected via the 250.29>125.00 and 265.35>170.00 transitions, respectively, recorded in multiple reaction monitoring (MRM) mode.

### 4. Directed evolution of GrsB1 for L-Piz

#### 4.1. Active site mutations

Mutation of the specificity code residues in A-domains is very likely to influence the overall substrate specificity[5,6]. To change the specificity of GrsB1 from L-Pro to L-Piz, we first compared its specificity code to A-domains annotated to be specific for L-Piz (Figure 1b) and identified differences in five positions. Accordingly, mutants GrsB1_Q663F, GrsB1_I729V, GrsB1_H755A, GrsB1_H755S, GrsB1_V763Y, and GrsB1_V764A were produced as proteins and assayed for their ability to produce PhePiz-DKP in an *in vitro* assay (Figure S2). By challenging the mutants with equimolar amounts of L-Pro and L-Piz, we were able to record the ratio of PhePiz-DKP to PhePro-DKP as a measure for improvements in L-Piz specificity. Mutations resulting in an increased specificity for L-Piz were combined, the respective proteins produced, purified, and analysed in repeated rounds of directed evolution. Finally, we obtained mutant GrsB1-AYA (GrsB1_H755A_V763Y_V7634) with a 40-fold increase in Piz specificity over GrsB1 (Figure S2).

#### 4.2. Targeted libraries of second shell positions

To further improve the activity of GrsB1-AYA, we continued to mutate "second shell residues". These residues are not directly in the active site but close. To find relevant differences, we analysed residues in a 5 Å sphere around the active site of GrsB1-AYA and compared them with the corresponding residues in different A-domains chosen to represent a broad array of different substrate specificities (Figure S1). It was found that all L-Piz activating A-domains showed conserved differences at positions 73 (T730L or T730F), 758 (P758Q), 761 (T761S), and 762 (H762F). To test the effect of these mutations, we created a library using degenerate codons in all four positions (730_HYT, 758_CMG, 761_WCT, 762_YWT). These codons included the respective residue of GrsB1-AYA, all identities observed in the Piz-activating A-domains, as well as some chemically similar residues resulting in a total library size of 96 possible mutants. This library was then screened using the lysate screening assay in 96 well format and candidates were ranked according to PhePiz-DKP yield and L-Piz specificity (Figure S3). The best candidates were sequenced, the respective proteins purified, and measured again in an *in vitro* DKP assay. This showed that the best candidate GrsB1-SQSF (GrsB1-AYA_T730S_P758Q_T761S_H762F) completely lost its activity for L-Pro and was exclusively active for L-Piz.

#### 4.3. Saturation mutagenesis of second shell positions

While GrsB1-SQSF showed exclusive specificity for L-Piz, its activity was strongly reduced compared to GrsB1. In further rounds of evolution, we aimed to improve enzyme activity while maintaining perfect L-Piz specificity. We applied saturation mutagenesis[7] to positions in the second shell that appeared variable in a sequence alignment of A-domains with various specificities. In a first round, NNK libraries in five positions (L634, C661, Y662, F703, P704) were analysed. Only mutations in positions 634 and 703 positively influenced PhePiz-DKP production. For both positions, the best candidates were combined to yield GrsB1-SQSF-VM (GrsB1-SQSF_L634V_F703M [Figure S4]). In the next round of evolution, NNK libraries for eight positions were analysed (T654, F658, V660, Q663, E664, S702, I729, A731). For positions 660, 663, and 731, several candidates showed improved production levels of PhePiz-DKP (V660M, V660P, V660S, Q663F, Q663W, Q663Y, A731V, A731G) and were subsequently combined in one library to account for all potential combinations including the wild-type identity at each of the three positions. In this way, we created mutant GrsB1-MWG (GrsB1-SQSF-VM_V660M_Q663W_A731G), which retains perfect specificity for L-Piz while having similar enzyme activity to GrsB1 (Figure S5).

### 5. Bioanalytical methods

#### 5.1. MesG/ hydroxylamine spectrophotometric assays

MesG assays for determination of adenylation kinetics[8] were performed in 384 well plates in a reaction volume of 100 µl. In each well, 2 µM of GrsB1 variant were placed in reaction buffer (50 mM Tris pH 7.6, 5 mM $MgCl_2$, 100 µM 7-methylthioguanosine [MesG], 150 mM $NH_2OH$ [adjusted to pH 7.5], 5 mM ATP, 1 mM TCEP, 0.4 U/ml inorganic phosphatase (I1643, Sigma), 1 U/ml purine nucleoside phosphorylase [N8264, Sigma]). Absorption at 355 nm was measured using a Synergy H1 plate reader (BioTek). First, the absorption was measured for 10 min at 30 °C without substrate to remove any phosphate contaminations. After addition of the respective substrate, the change in absorption at 355 nm was measured continuously over 30 min in 5 s intervals at 30 °C for all wells. Reactions containing buffer without substrate were monitored to obtain a background rate which was subsequently subtracted. Each measurement was done as biological duplicate and the mean used for calculations. The initial reaction velocity (OD/min) was plotted against the substrate concentration and non-linearly fitted to the Michaelis-Menten equation (Equation 1) to derive $K_M$ and $k_{cat}$ values using RStudio 2022.12.0

.

$$\frac{v_0}{E_0} = k_{cat}\frac{[S]}{K_M + [S]} \qquad (1)$$

#### 5.2. Thermal shift assay

Thermal shift assays were performed on an Applied Biosystems StepOne Real-Time PCR System using SYPRO Orange (Thermo Fisher Scientific) as fluorescence dye. The assay was carried out using 2 µM enzymes and 0 – 6.25 µM Pro-AMS or 0 – 50 µM Nip-AMS in 50 mM HEPES, 100 mM NaCl and 1 mM $MgCl_2$ at pH 8. Pro-AMS and Nip-AMS were prepared as 10-fold concentrated stock solutions and SYPRO Orange dye was diluted from a 5,000-fold to a 25-fold concentrated stock solution. The assay was carried out in 20 µl volume after combining 13 µl of enzyme solution, 2 µl of the inhibitor and 5 µl of the fluorescence dye. As negative control, the inhibitor was replaced with buffer. Temperature was kept at 25 °C for 2 min, increased to 99 °C over 40 min in 1 % increments and maintained at 99 °C for 2 min. All measurements were performed with two biological replicates and three technical replicates each. Resulting melting curves were analysed and the respective melting points calculated after the Boltzmann method using Protein Thermal Shift Software v1.3 (Thermo Fisher Scientific). A shift of the melting point ($\Delta T_m$) was calculated as the difference of the melting point measured with and without inhibitor at a series of inhibitor concentrations. By plotting $\Delta T_m$ against inhibitor concentration using a hyperbolic binding model (Equation 2), $K_D$ values for Pro-AMS and Nip-AMS were calculated for each enzyme.

$$\Delta T_m = \Delta T_m, max\ \frac{[I]}{(K_D + [I])} \qquad (2)$$

#### 5.3. Protein structure determination

Purified GrsB1 was crystallised by sitting-drop vapour diffusion on MRC2 96-well crystallisation plates (SwissSci) with 0.3 µL protein and 0.3 µL precipitant solution drops dispensed by Oryx8 robot (Douglas Instruments). GrsB1 was diluted to 9 mg/mL concentration in 20 mM HEPES buffer pH 7.5, 150 mM NaCl prior to dispensing. Initial crystals were obtained using Classic I 96-HTS (Jena Biosciences). Optimised crystals for data collection were grown in 20% w/v PEG 4000 and 312 mM sodium acetate solution. Crystals were soaked in cryoprotectant solution consisting of 20% PEG 4000 w/v, 312 mM sodium acetate and 30% w/v glycerol before flash-cooling in liquid nitrogen. X-ray data sets were recorded on the 10SA (PX II) beamline at the Paul Scherrer Institute (Villigen, Switzerland) at a wavelength of 1.0 Å using a Dectris Eiger3 16M detector with the crystals maintained at 100 K by a cryocooler. Diffraction data were integrated using XDS[9] and scaled and merged using AIMLESS[10]; data collection statistics are summarized in Table S3. The structure solution was automatically obtained by molecular replacement using pdb 5n81[3] as template. The map was of sufficient quality to enable 90% of the residues expected for the four copies of GrsB1 to be automatically fitted using Phenix autobuild.[11] The model was finalized by manual rebuilding in COOT[12] and refined using Phenix refine.[11]

### 6. Computational methods

#### 6.1. Docking simulations

Ligands and cofactors were docked into the active site of GrsB1 and GrsB1 mutant MWG using AutoDock Vina.[13,14] Homology model of GrsB1-MWG was created by YASARA Structure[15,16] using the solved structure of GrsB1 as template. Ligands were prepared for docking using Chimera Dock Prep to add hydrogens and charges.[17] When assessing the results, we selected ligand orientations in which the α-amino group of Pro or Piz was in close proximity to the conserved Asp659 of GrsB1; this orientation was not always the lowest possible energy solution. Initial docking results were then energy minimized using YASARA Structure. Figures of docking results were generated using PyMol.[18].

### 7. Production and isolation of $^{Piz}$GS

#### 7.1. Small scale optimisation of $^{Piz}$GS production conditions

Optimisation of production conditions was based on previously described conditions for heterologous production of GS in *E. coli*[4]. The Influence of different L-Piz concentrations in the medium as well as the effect of different cultivation media on the $^{Piz}$GS production were analysed. For small-scale production of $^{Piz}$GS, *E. coli* HM0079::pSU18-GrsB$_{Piz}$ was precultured in TB + cam medium over night at 37 °C and 230 rpm. The preculture was used to inoculate 5 ml of the respective culture medium + cam medium in a 100 mL Erlenmeyer flask at a starting $OD_{600}$ of 0.01. 6 mM L-Orn and the respective L-Piz concentration were added and the culture incubated at 30 °C and 400 rpm for 5 days. The cells were harvested in 2 ml Eppendorf tubes at 20,000 g for 5 min and the pellet resuspended in 1 ml 70% (v/v) EtOH and incubated for 2.5 h at 25 °C and 1000 rpm. Afterwards, the cell suspension was lysed in an ultrasonic bath for 15 min and the lysate cleared by centrifugation at 20,000 g for 5 min. The cleared lysate was diluted 1:20 (v/v) with 40% MeOH + 0.1% formic acid and analysed by UPLC-MS/MS.

To analyse the influence of L-Piz concentrations on $^{Piz}$GS production, 5 mM, 10 mM and 15 mM L-Piz were used in TB media following the procedure described above (Figure S12a).

For optimisation of culture conditions four different media were analysed: TB, LB, 2xYT and YPG. The previous results showed that 15 mM L-Piz resulted in the highest amount of $^{Piz}$GS produced and therefor this concentration was used for media comparison studies. The above described procedure was followed and the precultures were prepared in the respective media each (Figure S12b).

#### 7.2. Large scale production and purification of $^{Piz}$GS

For production of $^{Piz}$GS, *E. coli* HM0079::pSU18-GrsB$_{Piz}$ was precultured in LB + cam medium over night at 37 °C and 230 rpm. The preculture was used to inoculate 50 ml LB + cam medium in a 2 L Erlenmeyer flask at a starting $OD_{600}$ of 0.01. 6 mM L-Orn and 15 mM L-Piz were added and the culture incubated at 30 °C and 400 rpm for 5 days. Cells were separated from culture media by centrifugation at 10,000 g for 5 min. The cell pellet was resuspended in 70 % (v/v) EtOH and incubated at 25 °C and 400 rpm for 2.5 h. Afterwards, the cell suspension was lysed in an ultrasonic bath for 15 min and the lysate cleared by centrifugation at 20,000 g for 5 min. The cleared lysate was dried under vacuum. $^{Piz}$GS was extracted from the culture through repeated washes with petrol ether containing 0.2% (v/v) DIPEA as a base. The $^{Piz}$GS-containing wash fractions were combined and dried under vacuum. The residues of cell lysate as well as media extracts were redissolved in 40 % MeOH and applied to a Chromabond $C_{18}$ ec SPE column (Macherey-Nagel) preequilibrated with 40 % MeOH. The column was washed with 5 column volumes (CVs) each of 80 % MeOH and 100 % MeOH. $^{Piz}$GS was eluted with MeOH + 0.2% (v/v) DIPEA in 5 CV fractions. Product containing fractions were dried under vacuum. The residue was redissolved in 40 % (v/v) MeCN + 0.1% (v/v) trifluoroacetic acid (TFA) and submitted to semipreparative HPLC purification using a Shimadzu system with *dd*$H_2O$ + 0.1% TFA (A) and MeCN + 0.1% TFA (B) as solvents and a LunaC18 column (5 µm, 250 x 10 mm). Sample was purified using a gradient of 40-100% B over 17 min with a flow-rate of 5 ml/min. Fractions containing $^{Piz}$GS were combined and the solvent removed by freeze drying. Identity of the product was confirmed by UPLC-MS/MS measurement in MRM mode specific for $^{Piz}$GS (586>85), as well as HRMS (calculated: m/z = 586.3712 $[M+2H]^{2+}$, measured: m/z = 586.3713 $[M+2H]^{2+}$; Figure 3c).

### 8. Organic synthesis

#### 8.1. Analytics

NMR spectra were recorded in deuterated solvents (Carl Roth, Germany) on a Bruker AVANCE II 300 or Bruker AVANCE III 500 MHz spectrometer, equipped with a Bruker Cryoplatform. The chemical shifts (δ) are reported in parts per million (ppm) relative to the solvent residual peak of DMSO-d6 (1H: 2.50 ppm, quintet; 13C: 39.5 ppm, heptet). All reagents used were reagent grade and used as supplied (2',3'-O-isopropylideneadenosin purchased from Fisher Scientific, Cbz-L-Piz-OH purchased from BLD Pharmatech, Boc-D-Nip-OH purchased from Chempur, Boc-L-Pro-OSu purchased from Bachem). Sulfamoyl-isopropylideneadenosine was prepared as previously described[2]. Reactions were performed at ambient temperature under argon atmosphere in anhydrous solvents (Acros Organics) unless otherwise stated. Analytical thin-layer chromatography was performed on silica 60 F254 plates (0.25 mm, Merck). Compounds were visualized by dipping the plates in a ninhydrin/acetic acid solution followed by heating. Semipreparative HPLC purification was done on a Shimadzu system using *dd*$H_2O$ + 0.1% TFA (A) and MeCN + 0.1% TFA (B) as solvents and a Luna C18 column (5 µm, 250 x 10 mm).

#### 8.2. Synthesis of L-Piz

Cbz-L-Piz-OH (**1**, 210 mg, 0.8 mmol) and 10% palladium on carbon (66 mg, 2.72 mmol) were dissolved in MeOH (3.75 ml) under argon atmosphere and TFA (0.6 ml, 7.8 mmol) added. Triethylsilane (1.2 ml, 7.4 mmol) was added dropwise to prevent excessive gas evolution and the reaction stirred for 2.5 h at room temperature. Afterwards, the reaction was filtered over celite and volatiles removed under vacuum to yield L-Piz as TFA salt **2** (197 mg, 0.8 mmol, quantitative yield) as light-yellow crystals.

**$^1$H NMR** (300 MHz, DMSO) δ 3.75 (dd, *J* = 9.9, 3.3 Hz, 1H), 3.19 – 3.05 (m, 1H), 3.00 – 2.85 (m, 1H), 2.04 – 1.89 (m, 1H), 1.86 – 1.71 (m, 2H), 1.69 – 1.52 (m, 1H)**.**

**$^{13}$C NMR** (75 MHz, DMSO) δ 171.9, 56.3, 44.4, 25.4, 20.6.

#### 8.3. Synthesis of Boc-D-Nip-OSu

Boc-D-Nip-OH (**3**, 1 g, 4.4 mmol), *N*-hydroxysuccinimide (510 mg, 4.4 mmol) and *N*,*N*'-dicyclohexylcarbodiimide (910 mg, 4.4 mmol) were placed in a Schlenk tube under argon atmosphere. Dry THF (23 ml) was added and the reaction stirred at 0 °C for 30 min. Afterwards, the reaction was allowed to warm to room temperature and stirred for 18 h. The reaction was filtered and volatiles removed under vacuum from the filtrate to yield crude Boc-D-Nip-OSu **4** (1.77 g) that was used for the next steps without further purification.

**$^1$H NMR** (300 MHz, DMSO): δ 3.86 – 3.74 (m, 1H), 3.69 – 3.54 (m, 1H), 3.42 – 3.34 (m, 1H), 3.12 – 2.98 (m, 1H), 2.97 – 2.86 (m, 1H), 2.82 (s, 4H), 2.08 – 1.96 (m, 1H), 1.79 – 1.57 (m, 3H), 1.40 (s, 9H).

**$^{13}$C NMR** (75 MHz, DMSO) δ 170.0, 168.7, 79.0, 37.9, 27.9, 26.3, 25.4.

**HPLC-MS:** *m/z* = 327.80 $[M+H]^+$

#### 8.4. Synthesis of L-Pro-AMS

Sulfamoyl-isopropylideneadenosine (**5**, 350 mg, 0.91 mmol), Boc-L-Pro-OSu (**6**, 540 mg, 1.73 mmol) and $Cs_2CO_3$ (525 mg, 1.61 mmol) were dissolved in dry DCM (15 ml) under argon atmosphere. The reaction was stirred at room temperature for 16 h. Afterwards, the reaction was filtered, volatiles were removed under vacuum from the filtrate and the residues purified by column chromatography (silica 60, DCM/MeOH 85:15). Resulting Boc-L-Pro-AMS was further purified by semipreparative HPLC using a gradient from 5-95% B over 23 min with a flow rate of 8 ml/min. Product containing fractions were combined and volatiles removed under vacuum. 5 ml TFA/H2O 5:1 was added and the reaction stirred for 2 h at room temperature. Volatiles were removed under vacuum and the remaining reddish residue redissolved in $Et_2O$ upon which precipitate formed. The $Et_2O$ was decanted and the remaining precipitate dried under vacuum. The $Et_2O$ wash was repeated two times and each time the solid residue was ground intensely for 5 min with a spatula before decanting the $Et_2O$. The precipitate was dried and further purified by semipreparative HPLC using a gradient from 2-15% B over 10 min with a flow-rate of 8 ml/min. Product containing fractions were pooled and the sample freeze dried to yield L-Pro-AMS **7** (37 mg, 0.08 mmol) with 10% yield over two steps.

**$^1$H NMR** (500 MHz, DMSO): δ 8.59 (s, 1H), 8.37 (s, 1H), 5.95 (d, *J* = 5.6 Hz, 1H), 4.56 (t, *J* = 5.2 Hz, 1H), 4.19 – 4.16 (m, 2H), 4.16 – 4.12 (m, 2H), 3.95 – 3.90 (m, 1H), 3.25 – 3.15 (m, 1H), 3.15 – 3.04 (m, 1H), 2.21 – 2.12 (m, 1H), 1.95 – 1.87 (m, 1H), 1.87 – 1.75 (m, 2H).

**$^{13}$C NMR** (126 MHz, DMSO): δ 171.44 (s), 156.62 (s), 149.28 (s), 148.90 (s), 139.32 (s), 118.75 (s), 87.46 (s), 82.65 (s), 73.85 (s), 70.58 (s), 67.95 (s), 61.74 (s), 45.40 (s), 29.13 (s), 23.33 (s).

**HPLC-MS:** *m/z* = 444.1 $[M+H]^+$, *m/z* = 222.5 $[M+2H]^{2+}$

#### 8.5. Synthesis of D-Nip-AMS

**5** (150 mg, 0.39 mmol), **4** (243 mg, 0.74 mmol) and $Cs_2CO_3$ (225 mg, 0.69 mmol) were dissolved in dry DCM (8 ml) under argon atmosphere. The reaction was stirred at room temperature for 16 h. Afterwards, the reaction was filtered and volatiles were removed under vacuum. 4 N HCl in 1,4-dioxane (15 ml) was added and the reaction stirred for 2 h at room temperature. Volatiles were removed under vacuum and the crude residue was washed with DCM (3 x 10 ml). The residue was further purified by HPLC using a gradient from 2-15% B over 10 min with a flow rate of 8 ml/min. Product containing fractions were pooled and the sample freeze dried to yield D-Nip-AMS **8** (97 mg, 0.21 mmol) with 54% yield over two steps.

**$^1$H NMR** (500 MHz, DMSO): δ 8.69 (s, 1H), 8.46 (s, 1H), 6.41 (s, 1H), 4.93 (dd, *J* = 14.3, 1.5 Hz, 1H), 4.70 (dt, *J* = 4.3, 2.2 Hz, 1H), 4.61 (dd, *J* = 14.3, 2.6 Hz, 1H), 4.30 – 4.27 (m, 1H), 3.97 (d, *J* = 5.4 Hz, 1H), 3.23 – 3.03 (m, 2H), 2.98 – 2.82 (m, 2H), 1.93 – 1.81 (m, 1H), 1.79 – 1.69 (m, 1H), 1.62 – 1.47 (m, 2H).

**$^{13}$C NMR** (126 MHz, DMSO) δ 156.6, 149.3, 139.3, 119.5, 93.2, 83.2, 75.4, 70.0, 58.2, 44.4, 43.1, 25.9, 21.0.

**HPLC-MS:** *m/z* = 458.0 $[M+H]^+$, *m/z* = 229.0 $[M+2H]^{2+}$

### 9. Supplementary tables

**Table S1**: Primer sequences used for cloning.

| Name | Sequence (5' - 3') |
|---|---|
| GrsB1_piz_AflII_fw | TTA GCC AGA TTC TTA AGA GAA AAA GGC |
| GrsB1_piz_BspDI_rev | TCA TCC GCT TTA TCG ATA ACA ACA GC |
| pTrc99a-GrsB1_V969KpnI_fw | CAA ACG CAA AAT ATG TGG TAC CTA CAA ATG AGC TGG AAG AAA AAT TGG |
| pTrc99a-GrsB1_V969_rev | ACA TAT TTT GCG TTT GTA TTC ACA ATCC |
| pTrc99a-GrsB1_BamHI_rev | TGA TGA GAT CTG GAT CCC CCG TTT ATA TAA TTA G |
| GrsB1_piz_KpnI_2_rev | CAG CTC ATT TGT AGG TAC CAC ATA TTT TGC GTT TGT ATT CAC AAT CC |
| GrsB1_piz_H765A_fw | GTA CAT TTA CAC AAT GCT TAT GGT CCA TCA GAA ACG CAT G |
| GrsB1_piz_H765S_fw | GTA CAT TTA CAC AAT TCT TAT GGT CCA TCA GAA ACG CAT G |
| GrsB1_piz_H765_rev | ATT GTG TAA ATG TAC GTT ATG TTC ATG C |
| GrsB1_piz_I728V_fw | CGT GAA ACA TAT TGT CAC AGC AGG AGA ACA ATT AGT AGT TAA C |
| GrsB1_piz_I728V_rev | AAT ATG TTT CAC GCA AGT TGG AAA ACG |
| GrsB1_piz_Q673F_fw | TTT GAC GTG TGT TAC TTC GAA ATT TTT TCG ACG CTC TTG TC |
| GrsB1_piz_Q673F_rev | GTA ACA CAC GTC AAA ACT GCA TGT TG |
| GrsB1_piz_V773Y_fw | CAT CAG AAA CGC ATT ATG TTA CCA CCT ATA CTA TTA ATC CTG AAG C |
| GrsB1_piz_V773Y_rev | ATG CGT TTC TGA TGG ACC ATA ATG |
| GrsB1_piz_V774A_fw | CAG AAA CGC ATG TTG CTA CCA CCT ATA CTA TTA ATC CTG AAG CTG |
| GrsB1_piz_V774A_rev | AAC ATG CGT TTC TGA TGG ACC |
| GrsB1_piz_V773Y_V774A_fw | CCA TCA GAA ACG CAT TAT GCT ACC ACC TAT ACT ATT AAT CCT GAA GCT G |
| AYA_H761F_fw | GGT CCA TCA GAA ACG TTC TAT GCT ACC ACC TAT ACT ATT AAT CCT GAA GC |
| AYA_H761A_fw | GGT CCA TCA GAA ACG GCA TAT GCT ACC ACC TAT ACT ATT AAT CCT GAA GC |
| AYA_H761_rev | CGT TTC TGA TGG ACC ATA AGC ATT GTG |
| AYA_P757Q_fw | CAC AAT GCT TAT GGT CAG TCA GAA ACG CAT TAT GCT ACC ACC |
| GrsB1-AYA-757_rev | ACC ATA AGC ATT GTG TAA ATG TAC GTT ATG |
| GrsB1_piz_I611_nnk_fw | GAC TTG TTT TAT ATT NNK TAT ACA TCA GGA ACA ACA GGT AAA CC |
| GrsB1_piz_633nnk_fw | CAC AAA AAC ATC GTT AAT NNK CTC CAT TTT ACT TTC GAG AAA ACA AAT ATC AAC TTT AG |
| GrsB1_piz_660nnk_fw | TGC AGT TTT GAC GTG NNK TAT CAA GAA ATT TTT TCG ACG CTC TTG TC |
| GrsB1_piz_660_rev | CAC GTC AAA ACT GCA TGT TGT ATA C |
| GrsB1_piz_661_nnk_fw | AGT TTT GAC GTG TGT NNK CAG GAA ATT TTT TCG ACG CTC TTG TCT G |
| GrsB1_piz_Y661V_fw | GTT TTG ACG TGT GTG TCC AAG AAA TTT TTT CGA CGC TC |
| GrsB1_piz_661_rev | ACA CAC GTC AAA ACT GCA TG |

| | |
|---|---|
| GrsB1_piz_662Y_fw | TTT GAC GTG TGT TAC TAC GAG ATT TTT TCG ACG CTC TTG TCT GG |
| GrsB1_piz_662F_fw | TTT GAC GTG TGT TAC TTC GAG ATT TTT TCG ACG CTC TTG TCT GG |
| GrsB1_piz_662nnk_fw | TTT GAC GTG TGT TAC NNK GAG ATT TTT TCG ACG CTC TTG TCT GG |
| GrsB1_piz_662_rev | GTA ACA CAC GTC AAA ACT GCA TGT TG |
| GrsB1_piz_663nnk_fw | GAC GTG TGT TAC CAA NNK ATA TTT TCG ACG CTC TTG TCT GGA G |
| GrsB1_piz_663_rev | TTG GTA ACA CAC GTC AAA ACT GCA TGT TG |
| GrsB1_piz_653nnk_fw | AAA GTA TTA CAG TAT NNK ACT TGC AGT TTT GAC GTG TGT TAC C |
| GrsB1_piz_653_rev | ATA CTG TAA TAC TTT GTC ACT AAA GTT GAT ATT TGT TTT CTC G |
| GrsB1_piz_P659_Y662_fw | ACA TGC AGT TTT GAC CCT TGT TAC TAT GAG ATT TTT TCG ACG CTC TTG TCT GG |
| GrsB1_piz_S659_W662_fw | ACA TGC AGT TTT GAC AGT TGT TAC TGG GAG ATT TTT TCG ACG CTC TTG TCT GG |
| GrsB1_piz_M659_F662_fw | ACA TGC AGT TTT GAC ATG TGT TAC TTC GAG ATT TTT TCG ACG CTC TTG TCT GG |
| GrsB1_PIz_V659_F662_fw | ACA TGC AGT TTT GAC GTG TGT TAC TTC GAG ATT TTT TCG ACG CTC TTG TCT GG |
| GrsB1_piz_659nnk_fw | ACA TGC AGT TTT GAC NNK TGC TAC CAA GAA ATT TTT TCG ACG CTC TTG |
| GrsB1_piz_659V_fw | ACA TGC AGT TTT GAC GTG TGC TAC CAA GAA ATT TTT TCG ACG CTC TTG |
| GrsB1_piz_659M_fw | ACA TGC AGT TTT GAC ATG TGC TAC CAA GAA ATT TTT TCG ACG CTC TTG |
| GrsB1_piz_659S_fw | ACA TGC AGT TTT GAC TCG TGC TAC CAA GAA ATT TTT TCG ACG CTC TTG |
| GrsB1_piz_659P_fw | ACA TGC AGT TTT GAC CCG TGC TAC CAA GAA ATT TTT TCG ACG CTC TTG |
| GrsB1_piz_659_rev | GTC AAA ACT GCA TGT TGT ATA CTG TAA TAC TTT GTC |
| GrsB1_piz_657nnk_fw | TAT ACA ACA TGC AGT NNK GAT GTG TGT TAC CAA GAA ATT TTT TCG ACG |
| GrsB1_piz_657_rev | ACT GCA TGT TGT ATA CTG TAA TAC TTT GTC AC |
| GrsB1_piz_701nnk_fw | AAT ATT GAA GTA TTA NNK TTA CCT GTG GCT TTT CTA AAA TTT ATT TTC AAT G |
| GrsB1_piz_701_rev | TAA TAC TTC AAT ATT TTC ACG TTT TAC TAA ATC AAA TAA TTG C |
| GrsB1_piz_702wtk_fw | ATT GAA GTA TTA TCC WTK CCC GTG GCT TTT CTA AAA TTT ATT TTC AAT GAA AG |
| GrsB1_piz_702_rev | GGA TAA TAC TTC AAT ATT TTC ACG TTT TAC TAA ATC |
| GrsB1_piz_703_nnk_fw | GAA GTA TTA TCC TTT NNK GTA GCT TTT CTA AAA TTT ATT TTC AAT GAA AGA GAA TTT ATC |
| GrsB1_piz_728nnk_fw | TGC GTG AAA CAT ATT NNK TCA GCA GGA GAA CAA TTA GTA GTT AAC AAT GAG |
| GrsB1_piz_728_rev | AAT ATG TTT CAC GCA AGT TGG AAA ACG |
| GrsB1-AYA-729HYT_fw | CGT TTT CCA ACT TGC GTG AAA CAT ATT ATC HYT GCA GGA GAA CAA TTA GTA GTT AAC AAT GAG |
| GrsB1-AYA-729_rev | GAT AAT ATG TTT CAC GCA AGT TGG AAA ACG |
| pTrc99a_GrsAB1_EcoNI_fw | ATT TCA CAC AGG AAA CTC GAG GCA GAA TAC CTA ACA AAG GAA TCG G |

| | |
|---|---|
| pTrc99a_GrsAB1_EcoNI_rev | TGG TGA TGA GAT CTG GAT CCT TGA ATG CCT AAC GTA GGA AGT TCC |
| GrsAB1_grsTAB_fw | TAC GCA GAA TAC CTA ACA AAG GAA TCG G |
| GrsAB1_grsTAB_rev | TTA TAT TGA ATG CCT AAC GTA GGA AGT TCC |
| pOPINF-GrsB1-A_3_fw | AAG TTC TGT TTC AGG GCC CGG ATT CGA TAA CAG AGT ATC CTG ATA AGA CG |
| pOPINF_GrsB1-A-NTsub_rev | ATG GTC TAG AAA GCT TTA GGC TCT TCC TAA AAA TTC GAT ATT TCC GTC |
| GrsB1_piz_730gbg_fw | AAA CAT ATT ATC TCT GBG GGT GAA CAA TTA GTA GTT AAC AAT GAG TTT AAA CG |
| GrsB1_piz_730_rev | AGA GAT AAT ATG TTT CAC GCA AGT TGG |

**Table S2**: Single mutations in each GrsB1 mutant generation.

| | **GrsB1-AYA** | **GrsB1-SQSF** | **GrsB1-SQSF-VM** | **GrsB1-MWG** |
|---|---|---|---|---|
| **First generation** | H755A | H755A | H755A | H755A |
| | V763Y | V763Y | V763Y | V763Y |
| | V764A | V764A | V764A | V764A |
| **Second generation** | | T730S | T730S | T730S |
| | | P758Q | P758Q | P758Q |
| | | T761S | T761S | T761S |
| | | H762F | H762F | H762F |
| **Third generation** | | | L634V | L634V |
| | | | F703M | F703M |
| **Fourth generation** | | | | V660M |
| | | | | Q663W |
| | | | | A731G |

**Table S3.** Data collection and refinement statistics.*

| | GrsB1 |
|---|---|
| **Wavelength** | |
| **Resolution range** | 59.9 - 2.6 (2.693 - 2.6) |
| **Space group** | R 3 2 :H |
| **Unit cell** | 195.021 195.021 346.46 90 90 120 |
| **Total reflections** | 816413 (83498) |
| **Unique reflections** | 77790 (7694) |
| **Multiplicity** | 10.5 (10.9) |
| **Completeness (%)** | 99.95 (100.00) |
| **Mean I/sigma(I)** | 12.86 (1.08) |
| **Wilson B-factor** | 68.33 |
| **R-merge** | 0.1426 (2.368) |
| **R-meas** | 0.15 (2.485) |
| **R-pim** | 0.04604 (0.752) |
| **CC1/2** | 0.999 (0.434) |
| **CC*** | 1 (0.778) |
| **Reflections used in refinement** | 77781 (7694) |
| **Reflections used for R-free** | 1990 (196) |
| **R-work** | 0.2228 (0.3365) |
| **R-free** | 0.2526 (0.3476) |
| **CC(work)** | 0.950 (0.612) |
| **CC(free)** | 0.950 (0.635) |
| **Number of non-hydrogen atoms** | 12893 |
| **macromolecules** | 12840 |
| **solvent** | 53 |
| **Protein residues** | 1560 |
| **RMS(bonds)** | 0.003 |
| **RMS(angles)** | 0.62 |

| | |
|---|---|
| **Ramachandran favored (%)** | 98.32 |
| **Ramachandran allowed (%)** | 1.68 |
| **Ramachandran outliers (%)** | 0.00 |
| **Rotamer outliers (%)** | 1.40 |
| **Clashscore** | 8.46 |
| **Average B-factor** | 74.72 |
| **macromolecules** | 74.76 |
| **solvent** | 65.74 |

*Statistics for the highest-resolution shell are shown in parentheses.

### 10. Supplementary figures

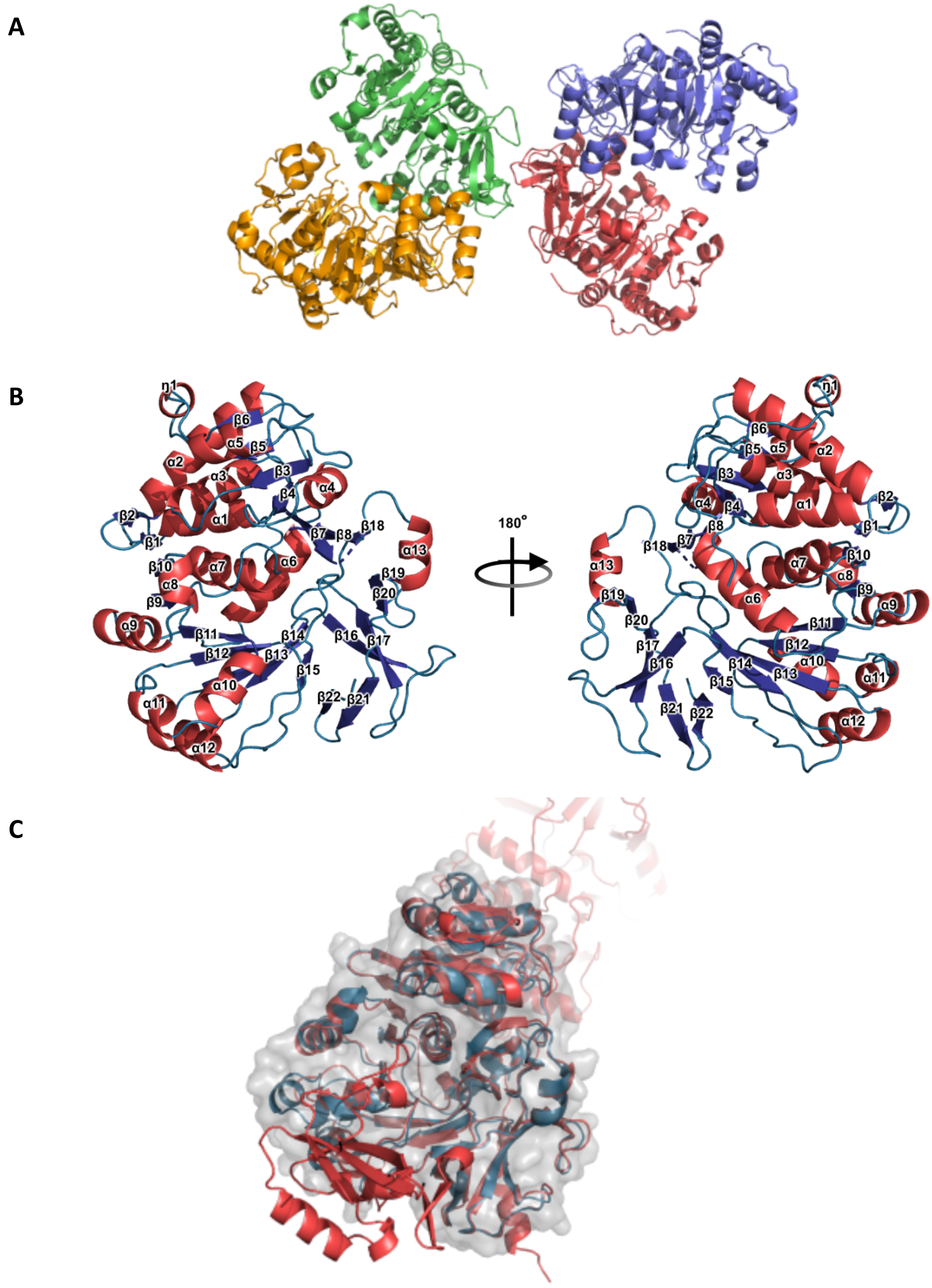

**Figure S1** Structure of GrsB1-$A_{core}$. **a)** Cartoon representation of GrsB1 crystallised as a homotetramer at 2.6 Å, coloured by chains. **b)** Cartoon of GrsB1 monomer coloured by secondary structure. **c)** Alignment of GrsB1 (blue) to GrsA-A (red, PDB 1AMU) with an RMSD of 0.820 Å.[19]

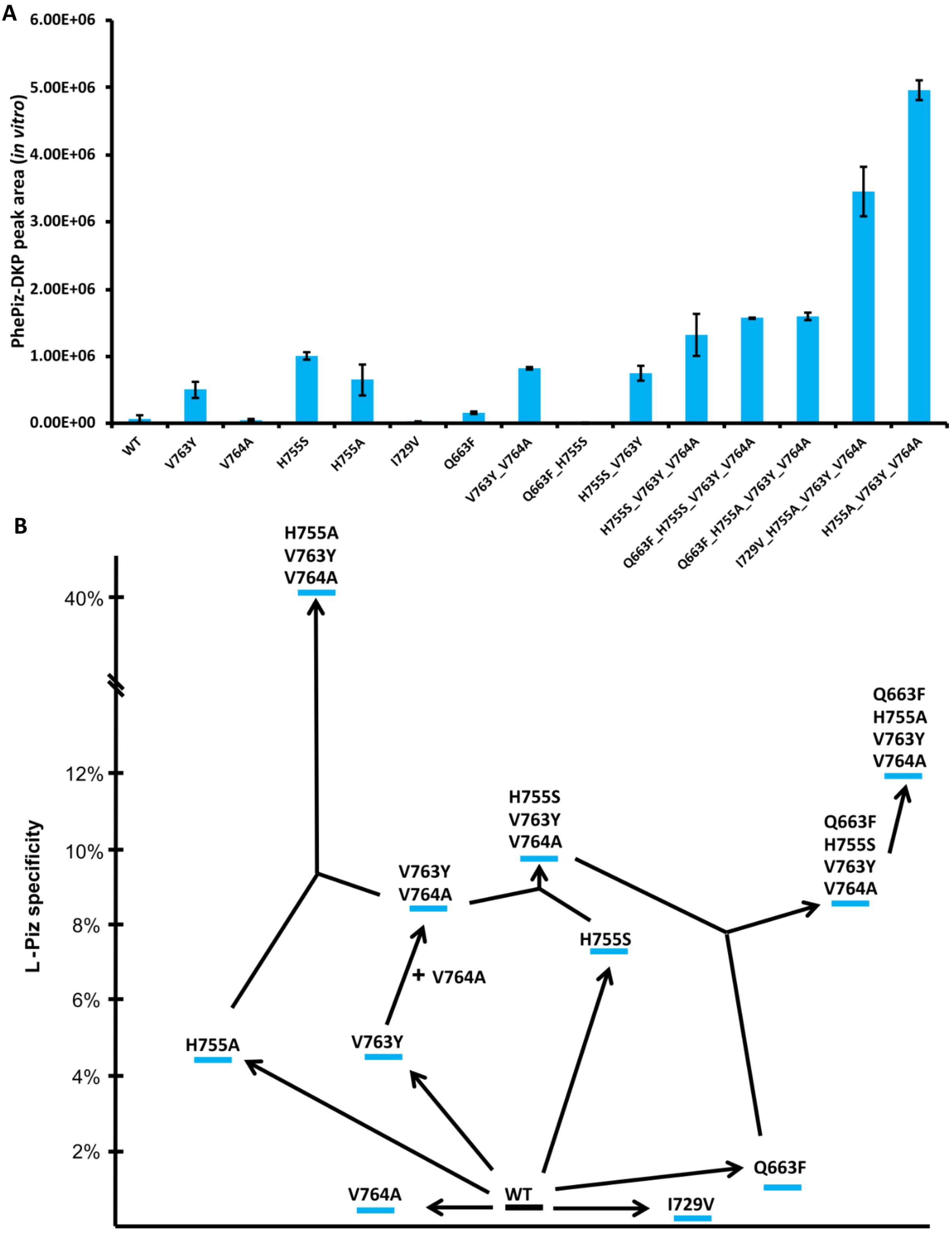

**Figure S2: a)** PhePiz-DKP peak areas of the 260>120 transition for purified GrsB1 active site mutants after the *in vitro* DKP assay measured by UPLC-MS/MS in MRM mode. The error bars indicate the standard deviation for technical triplicates. **b)** Evolutionary tree of L-Piz specificity for mutants obtained during GrsB1 active site mutagenesis. As a measure of specificity, the ratio of PhePiz-DKP and PhePro-DKP peak areas is used measured under substrate competition. Single mutants were characterized and beneficial mutations combined to obtain mutant GrsB1H755A_V763Y_V764A which will be named GrsB1-AYA from here on.

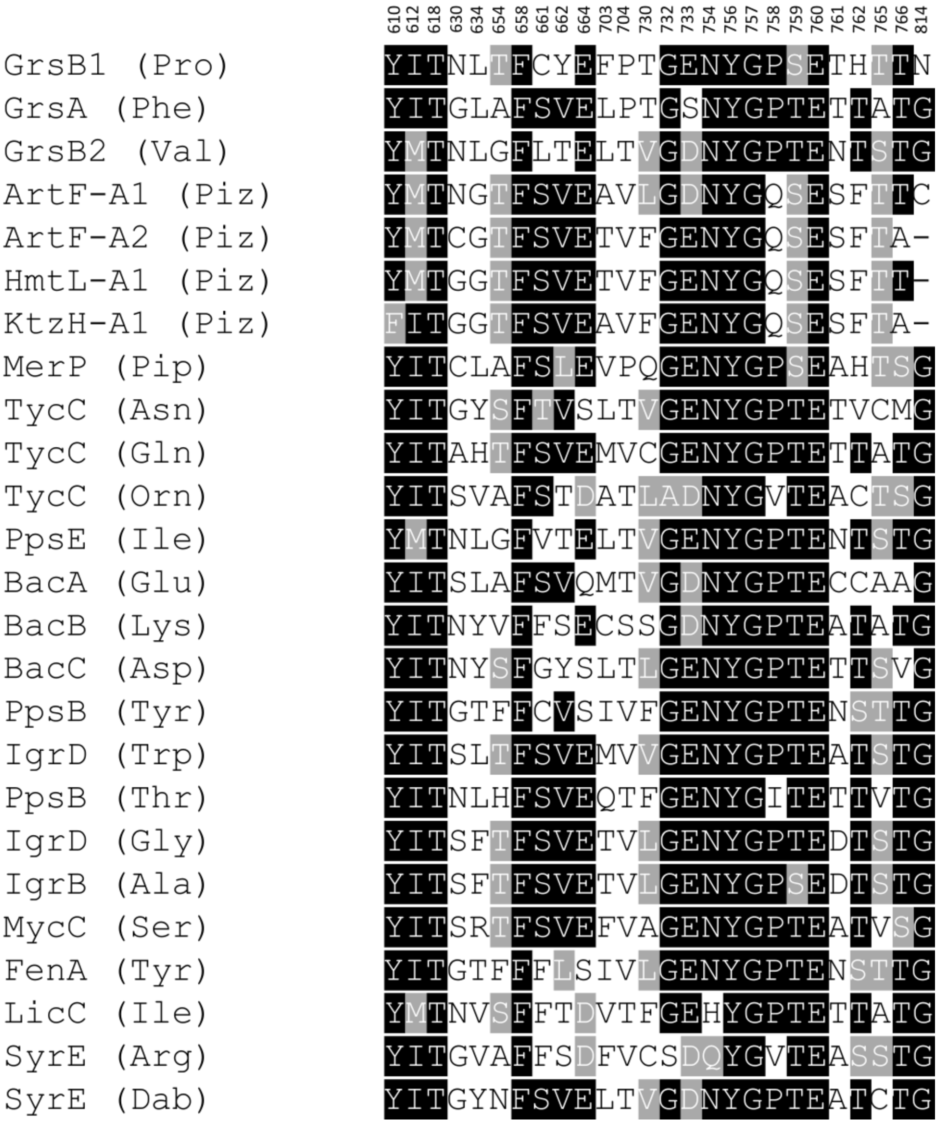

**Figure S3:** Comparison of second shell residues at 5 Å distance from the active site of GrsB1 and other A-domains selected to represent a broad array of different substrate specificities. Specificity code residues (first shell) are excluded.

A

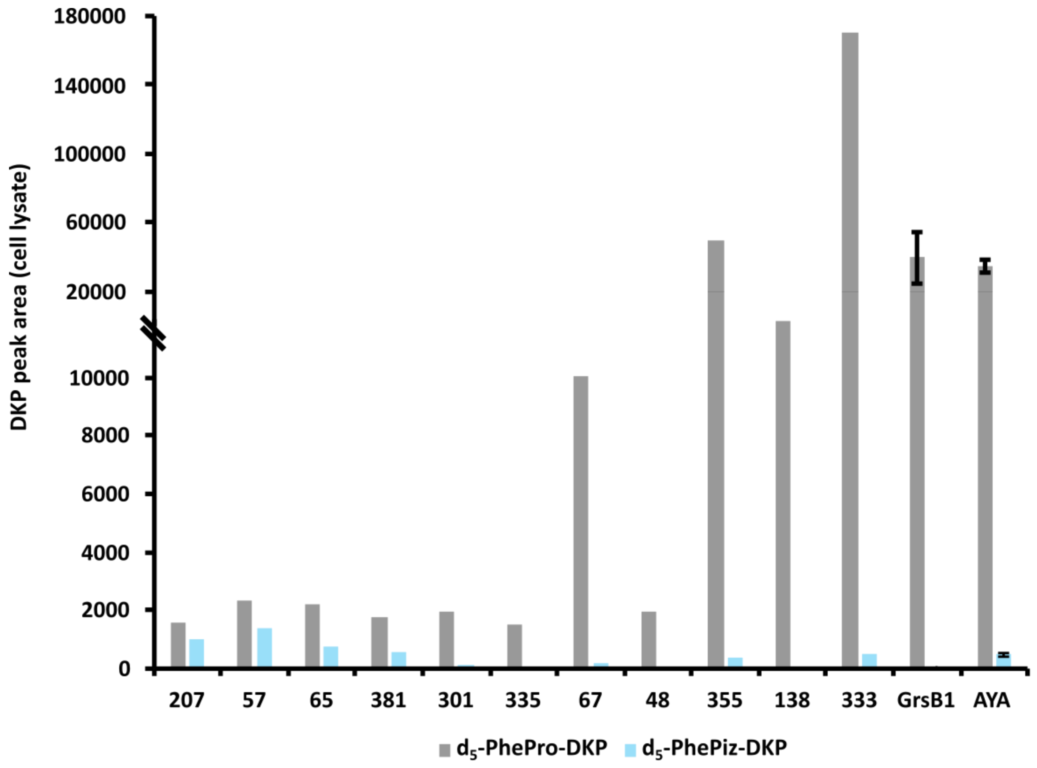

B

| | Mutations | L-Piz specificity | | Mutations | L-Piz specificity |
|---|---|---|---|---|---|
| **GrsB1** | - | 0.1% | **335** | AYA, T730F , P758Q, T761S, H762F | 4% |
| **AYA** | H755A, V763Y, V764A | 1.5% | **67** | AYA, T730L , H762F | 2% |
| **207** | AYA, T730S , P758Q, T761S, H762F | 63% | **48** | AYA, T730L , P758Q, H762L | 1% |
| **57** | AYA, T730S , P758Q, H762F | 60% | **355** | AYA, T730I , H762F | 0.8% |
| **65** | AYA, P758Q , H762F | 34% | **138** | AYA, T730S , P757Q, T761S | 0.3% |
| **381** | AYA, P758Q , H762Y | 30% | **333** | AYA, T730S , H762L | 0.3% |
| **301** | AYA, P758Q , T761S, H762L | 5% | | | |

**Figure S4**: Second generation of GrsB1 mutants. **a)** $d_5$-PhePro-DKP and $d_5$-PhePiz-DKP peak areas as measured during lysate screening. Peak areas were determined by UPLC-MS/MS in MRM mode following the 250>125 and 265>170 transitions, respectively. GrsB1 and GrsB1-AYA were analysed as controls under the same conditions. Peak areas for the controls are the mean of four biological replicates each. Error bars indicate the standard deviation. GrsB1-AYA shows a smaller signal for PhePiz-DKP when compared to *in vitro* assay with purified protein (Figure S2a) due to measuring the less intense 265>170 transition to prevent background signals from the cell lysate. **b)** L-Piz specificity for different mutations shown in comparison to GrsB1 and AYA. Mutant 207 was renamed as SQSF in following experiments.

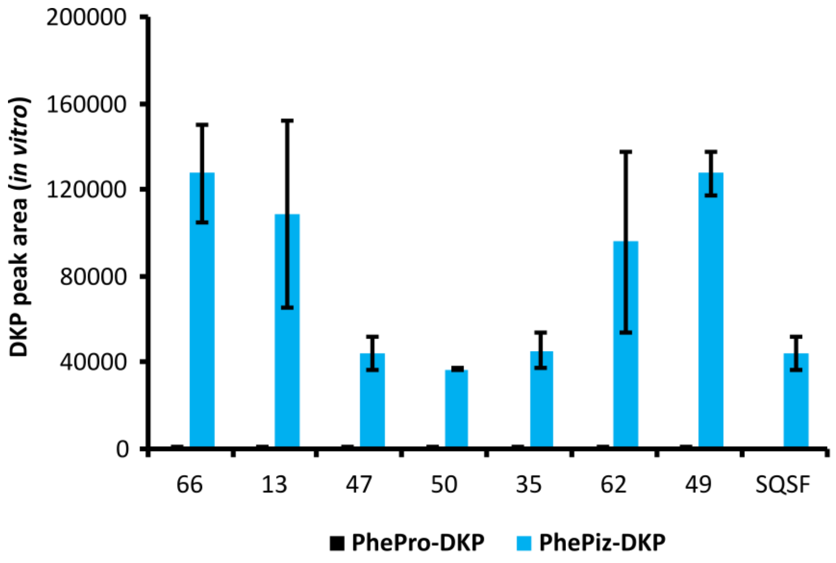

| | Mutations | | Mutations |
|---|---|---|---|
| **SQSF** | T730S, H755A, P758Q, T761S, H762F, V763Y, V764Y | **50** | SQSF, L634S |
| **66** | SQSF, L634V | **35** | SQSF, L634F |
| **13** | SQSF, L634A | **62** | SQSF, F703L |
| **47** | SQSF, L634G | **49** | SQSF, F703M |

**Figure S5:** Third generation of GrsB1 mutants. PhePro-DKP and PhePiz-DKP peak areas as measured in the *in vitro* DKP assay with purified proteins under substrate competition. Peak areas were determined by UPLC-MS/MS in MRM mode following the 240>120 and 260>120 transitions, respectively and represent the mean of two biological replicates. Error bars indicate the standard deviation. First, good candidates were identified by lysate screening. Then, the selected proteins were purified and used in an *in vitro* DKP formation assay. Purified GrsB1-SQSF was used as a control for the previous generation and measured in biological duplicates. Compared to the lysate screening (Figure S3), SQSF showed higher Piz/Pro ratios when measured with purified protein due to the better controlled conditions in the *in vitro* DKP assay allowing to measure the more intense 260>120 transition. Mutations 66 and 49 were combined and named SQSF-VM in following experiments.

A

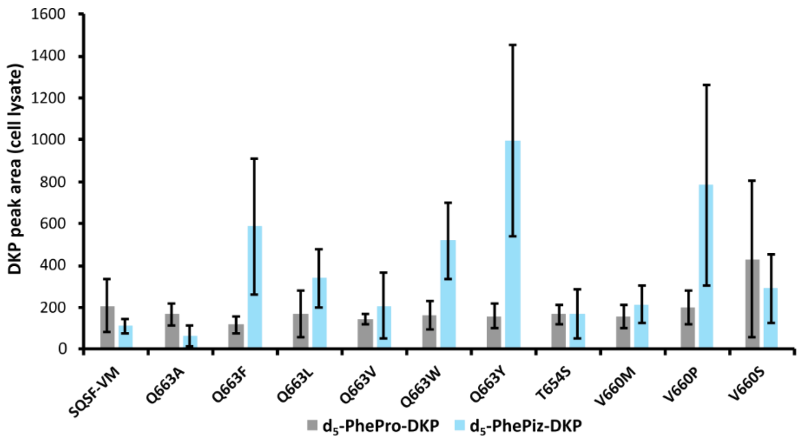

B

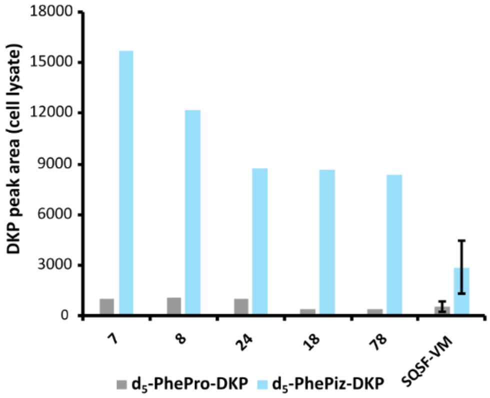

| | Mutations |
|---|---|
| **SQSF-VM** | L634V, F703M, T730S, H755A, P758Q, T761S, H762F, V763Y, V764A |
| **7** | SQSF-VM, A731G |
| **8** | SQSF-VM, A731A |
| **24** | SQSF-VM, A731G |
| **18** | SQSF-VM, A731A |
| **78** | SQSF-VM, A731A |

C

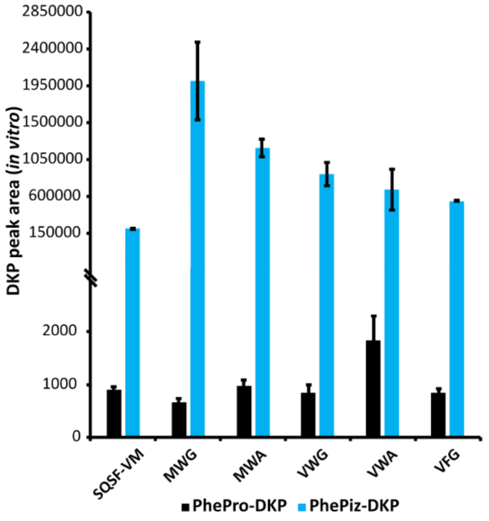

| | Mutations |
|---|---|
| **SQSF-VM** | L634V, F703M, T730S, H755A, P758Q, T761S, H762F, V763Y, V764A |
| **MWG** | SQSF-VM, V660M, Q663W, A731G |
| **MWA** | SQSF-VM, V660M, Q663W |
| **VWG** | SQSF-VM, Q663W, A731G |
| **VWA** | SQSF-VM, Q663W, |
| **VFG** | SQSF-VM, Q663F, A731G |

**Figure S6**: Fourth generation of GrsB1 mutants. **a)** Results of lysate screening for positions 654, 660 and 663. After individual screening of single plates for each position, the best candidates were all rescreened on the same plate. SQSF-VM was used as control. Shown are $d_5$-PhePro-DKP and $d_5$-PhePiz-DKP peak areas measured in MRM mode following the 250>125 and 265>170 mass transition. respectively. Peak areas represent the mean of six biological replicates and the error bars indicate the respective standard deviations. **b)** Results of lysate screening for positions 731. Shown are $d_5$-PhePro-DKP and $d_5$-PhePiz-DKP peak areas measured in MRM mode following the 250>125 and 265>170 mass transition, respectively, for the five best mutants. Peak areas for SQSF-VM as control represent the mean of four biological replicates. The error bars indicate the respective standard deviations. **c)** Results of *in vitro* DKP assays for mutants combining the best candidates for positions 660, 663 and 731. After initial lysate screening of the combination library, the most promising mutants were identified and the respective proteins purified. Purified SQSF-VM was used as a control. The shown peak areas represent the mean of two biological replicates. The error bars indicate the standard deviation.

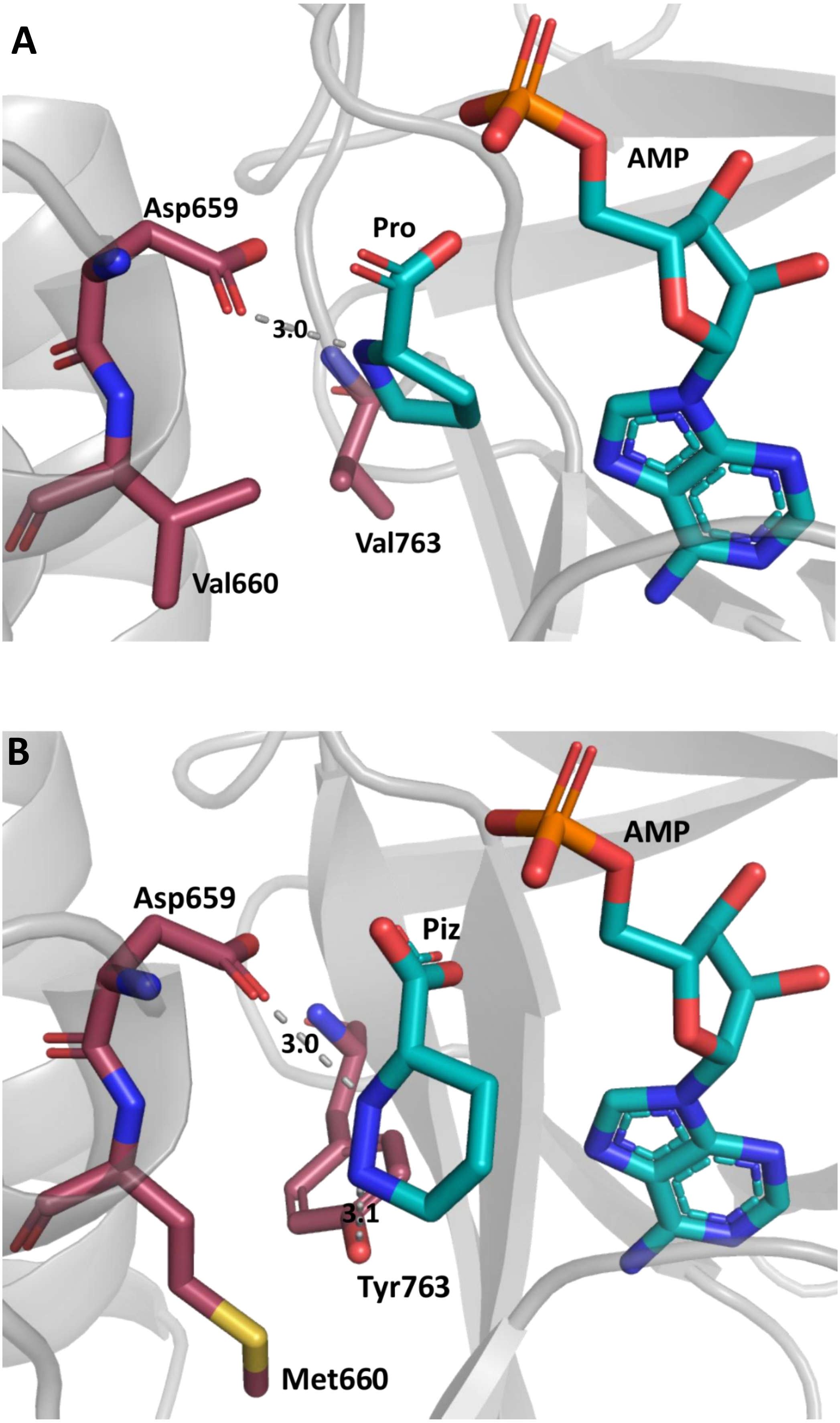

**Figure S7**: **a)** GrsB1 docked with substrate Pro and ligand AMP. Interaction between Asp659 and the α-amino group of Pro is indicated. **b)** Model of GrsB1-MWG docked with Piz and ligand AMP. Potential interactions between Asp659 and α-amino group of Piz and Tyr763 and the distal amino group of Piz are indicated.

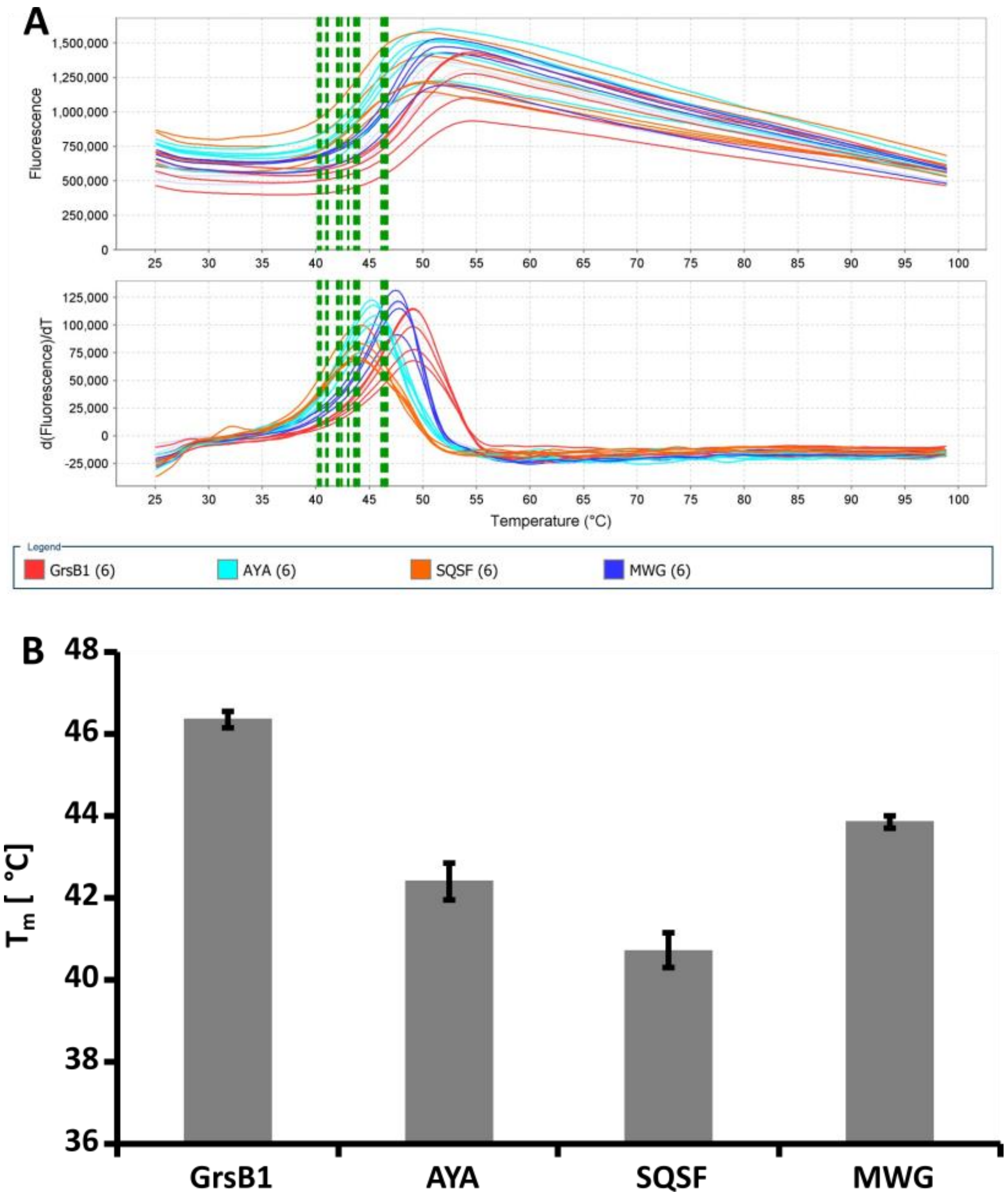

**Figure S8**: **a)** Melting curves from the thermal shift assay of GrsB1 and GrsB1 mutants AYA, SQSF and MWG to compare thermal stability between the different mutant generations. Derived melting points are indicated by black lines. **b)** Comparison of melting points of GrsB1 mutants. Error bars indicate standard deviations for two biological replicates measured as technical triplicate each.

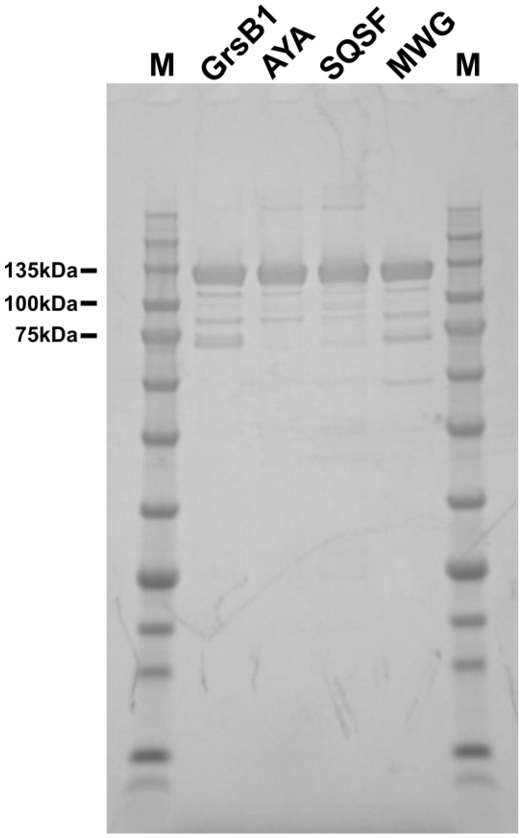

| Protein | Yield (mg/L) |
|---|---|
| GrsB1 | 51 ± 16 |
| AYA | 40 ± 22 |
| SQSF | 38 ± 3 |
| MWG | 62 ± 10 |

**Figure S9**: SDS PAGE of all relevant GrsB1 mutant generations after Ni affinity purification. 1 µg protein was loaded per well. Proteins were stained by Quick Coomassie stain (Serva). M: Triple Color Protein Standard III (Serva). Protein concentrations were calculated by measuring absorption at 280 nM using calculated extinction coefficients.

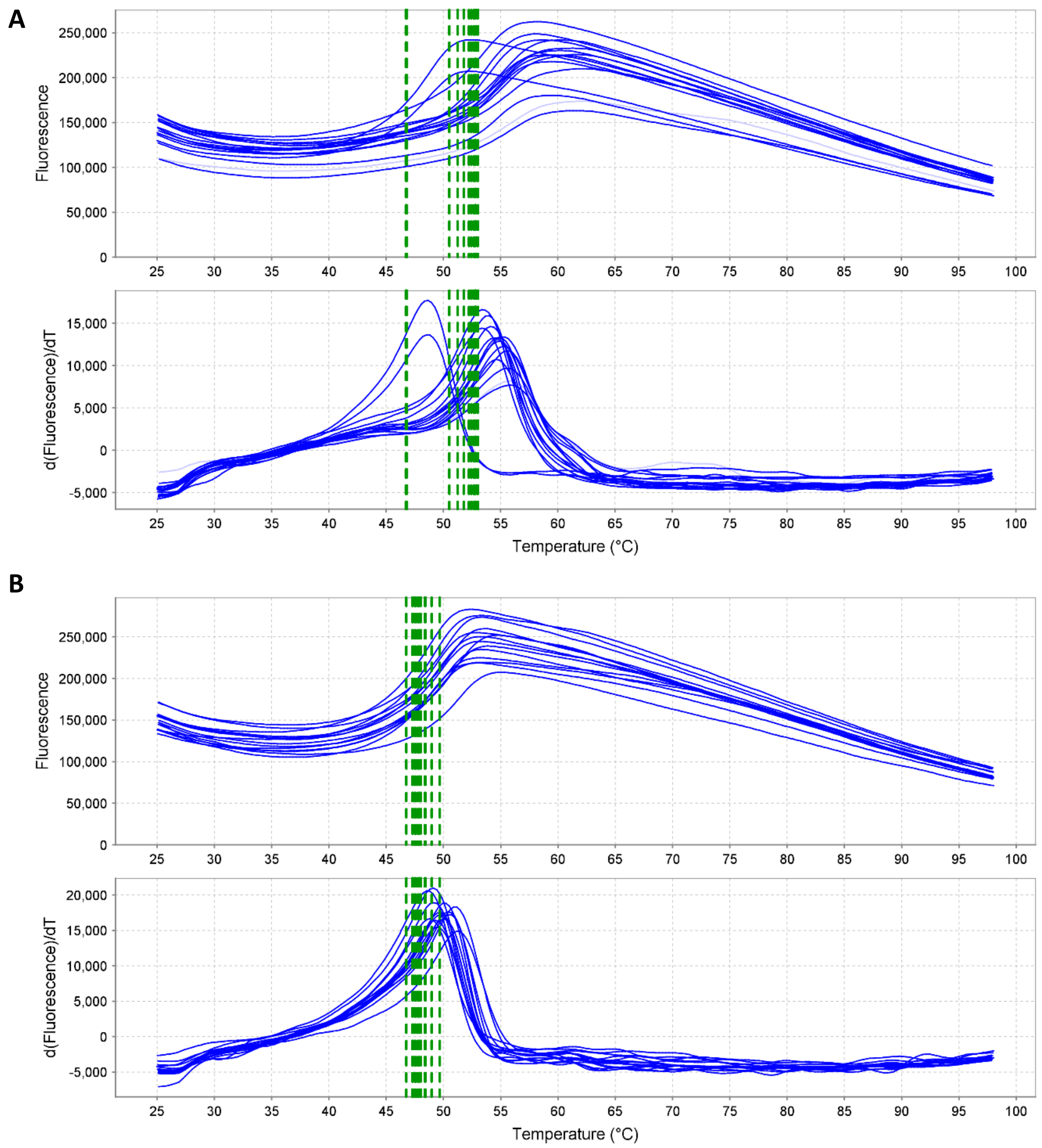

**Figure S10**: Melting curves from the thermal shift assay of GrsB1 titrated with inhibitors **a)** L-Pro-AMS and **b)** D-Nip-AMS. The calculated melting temperatures are indicated by the green line.

A

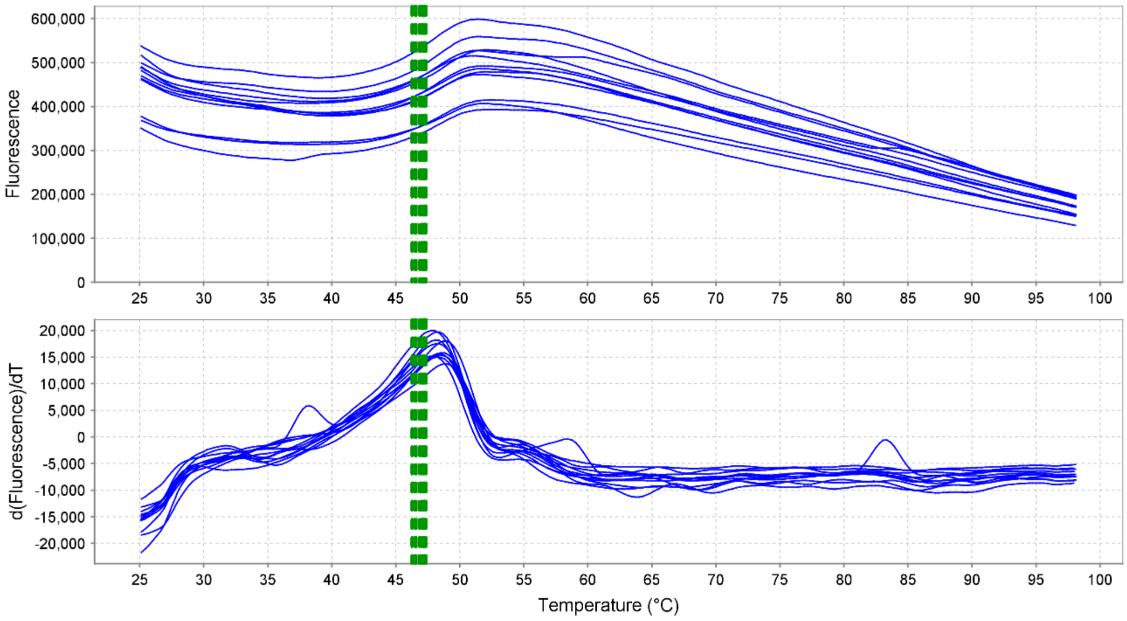

B

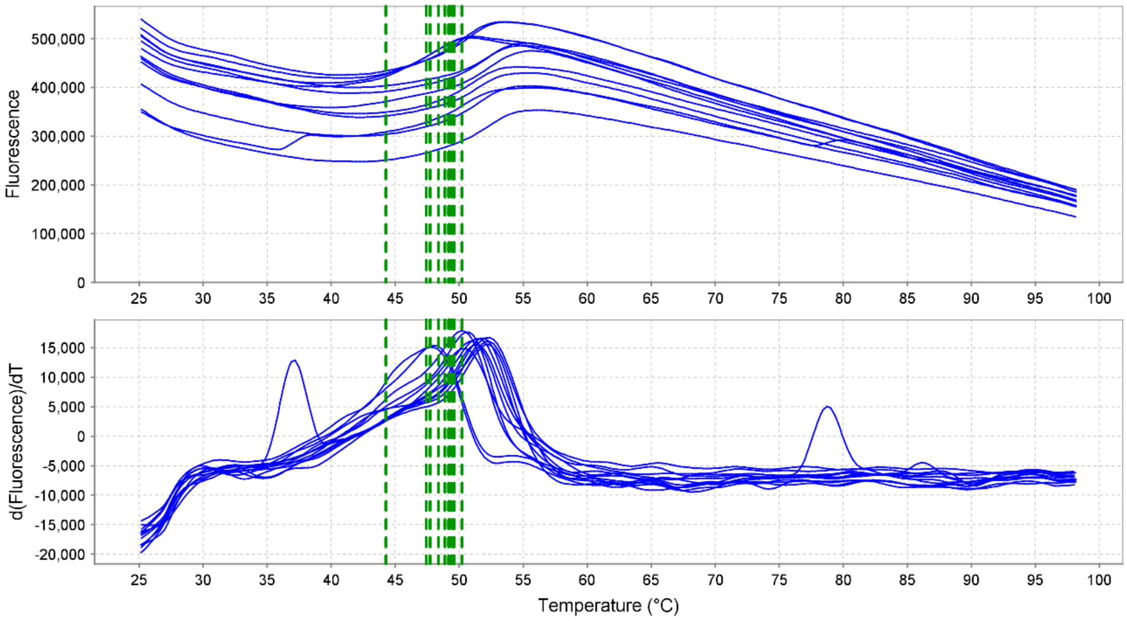

**Figure S11:** Melting curves from the thermal shift assay of MWG titrated with inhibitors **a)** L-Pro-AMS and **b)** D-Nip-AMS. The derived melting temperatures are indicated by the green line.

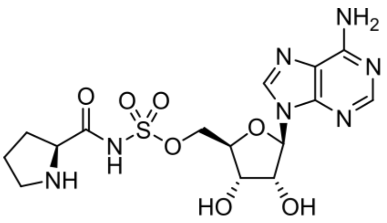

**Pro-AMS (7)**

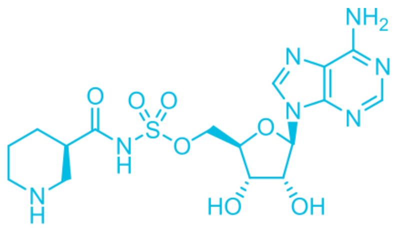

**Nip-AMS (8)**

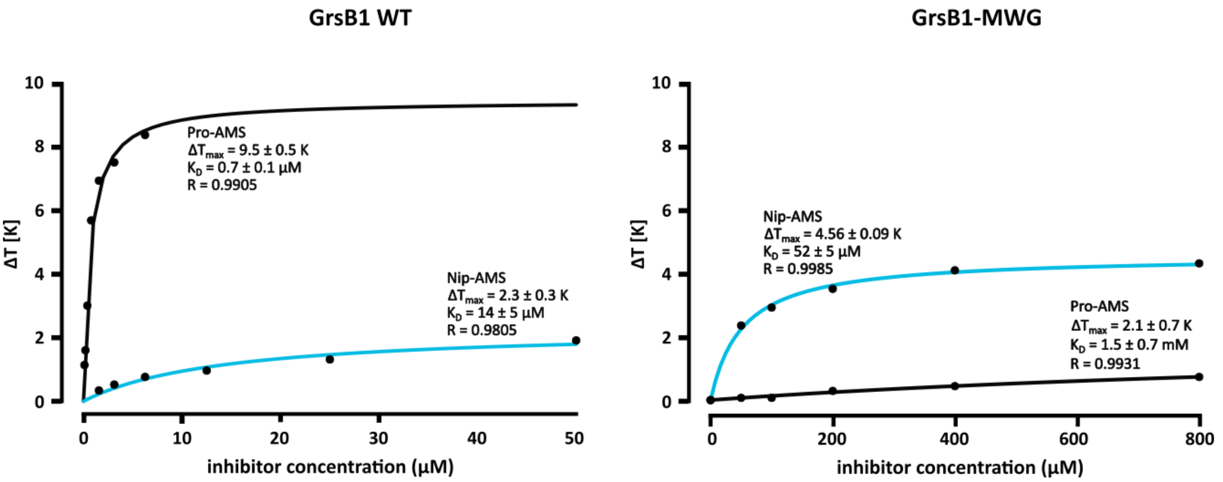

**Figure S12**: Shift in melting temperature of WT GrsB1 and GrsB1-MWG when titrated with L-Pro-AMS (**7**) and D-Nip-AMS (**8**). Melting points were measured using thermal shift assays and fitted using Equation 2 with the mean of two biological replicates. Calculated errors indicate error of the fit.

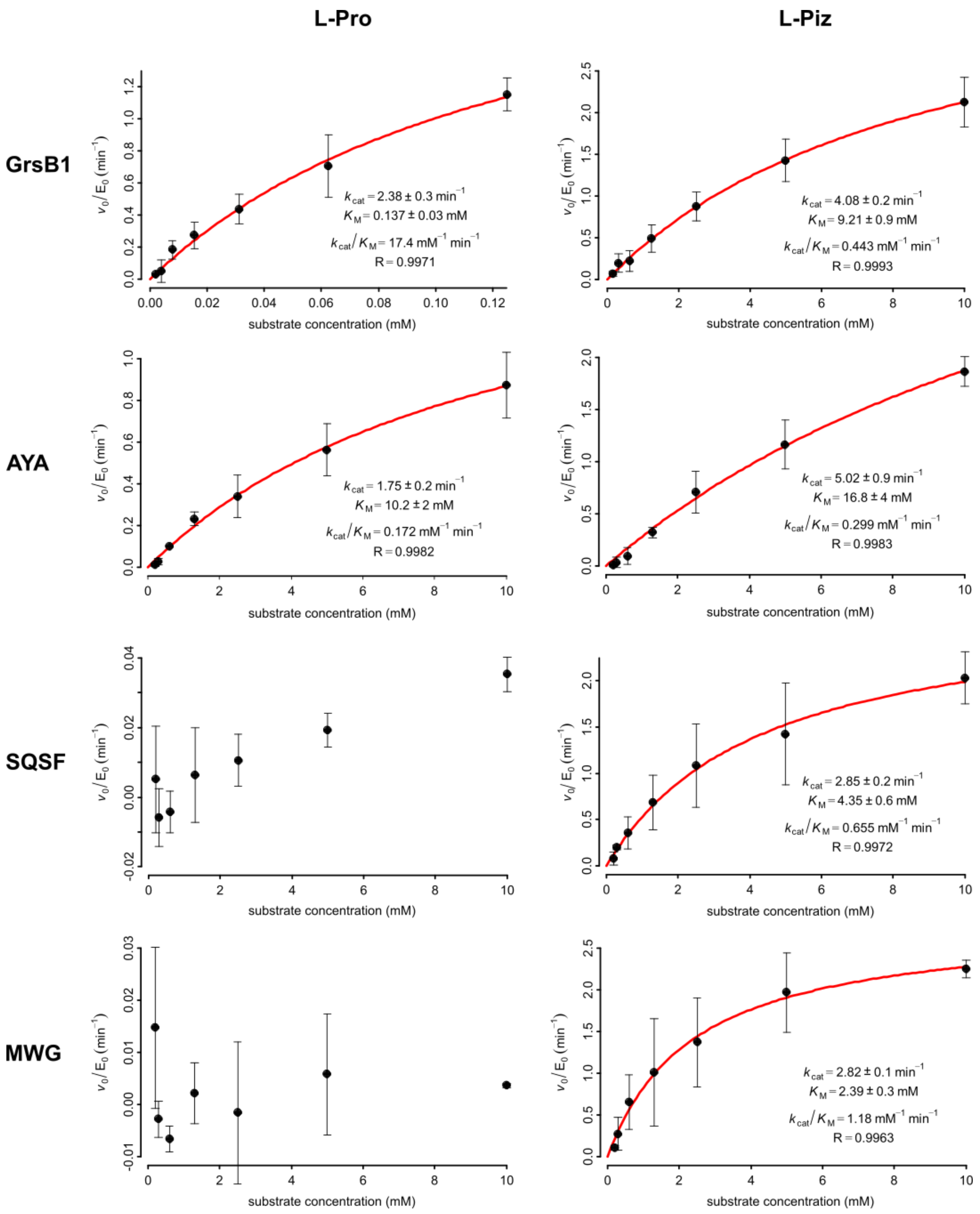

**Figure S13**: Adenylation kinetics of relevant GrsB1 mutants with L-Pro and L-Piz as substrate respectively. SQSF and MWG showed no detectable activity for L-Pro. Error bars indicate standard deviation of two biological replicates. Calculated errors indicate error of the fit when using the mean of two biological replicates.

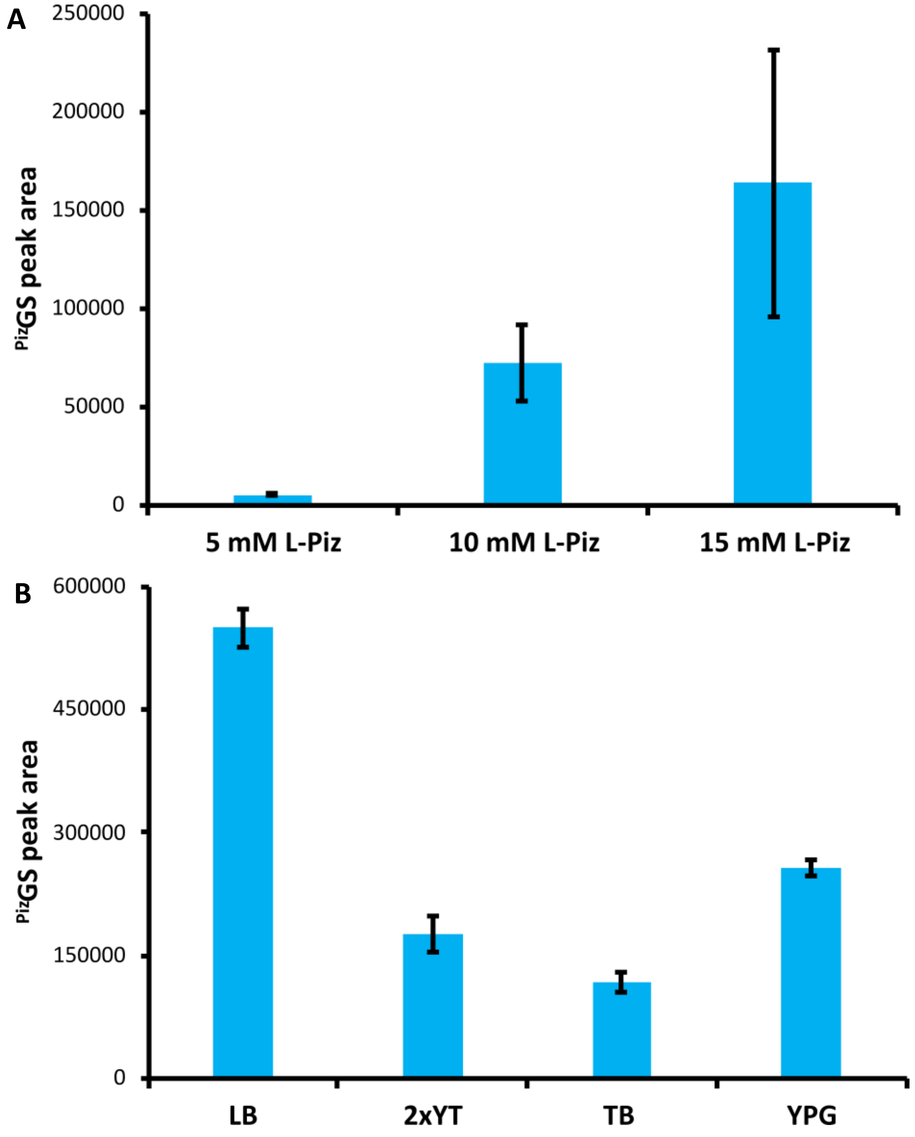

**Figure S14**: Optimisation of $^{Piz}$GS production in *E. coli*. Peak areas for $^{Piz}$GS were measured by UPLC-MS/MS in MRM mode following the 586>120 transition. Error bars indicate standard deviation of measurements for three biological replicates. **a)** Influence of different L-Piz concentrations on $^{Piz}$GS production when cultivated in TB medium. **b)** Influence of different cultivation media on $^{Piz}$GS production when adding 15 mM L-Piz to the medium.

### 11. NMR spectra

**L-Piperazic acid (2) $^1$H NMR**

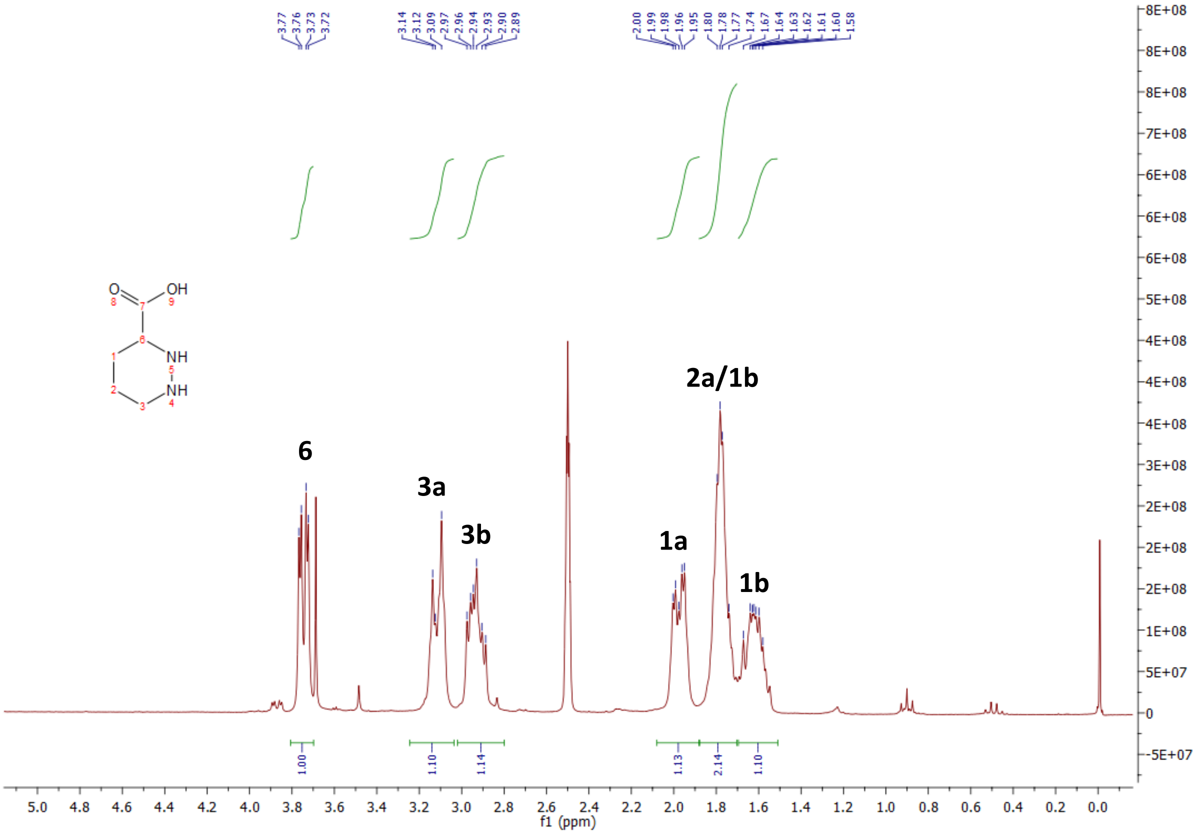

**L-Piperazic acid (2) $^{13}$C NMR**

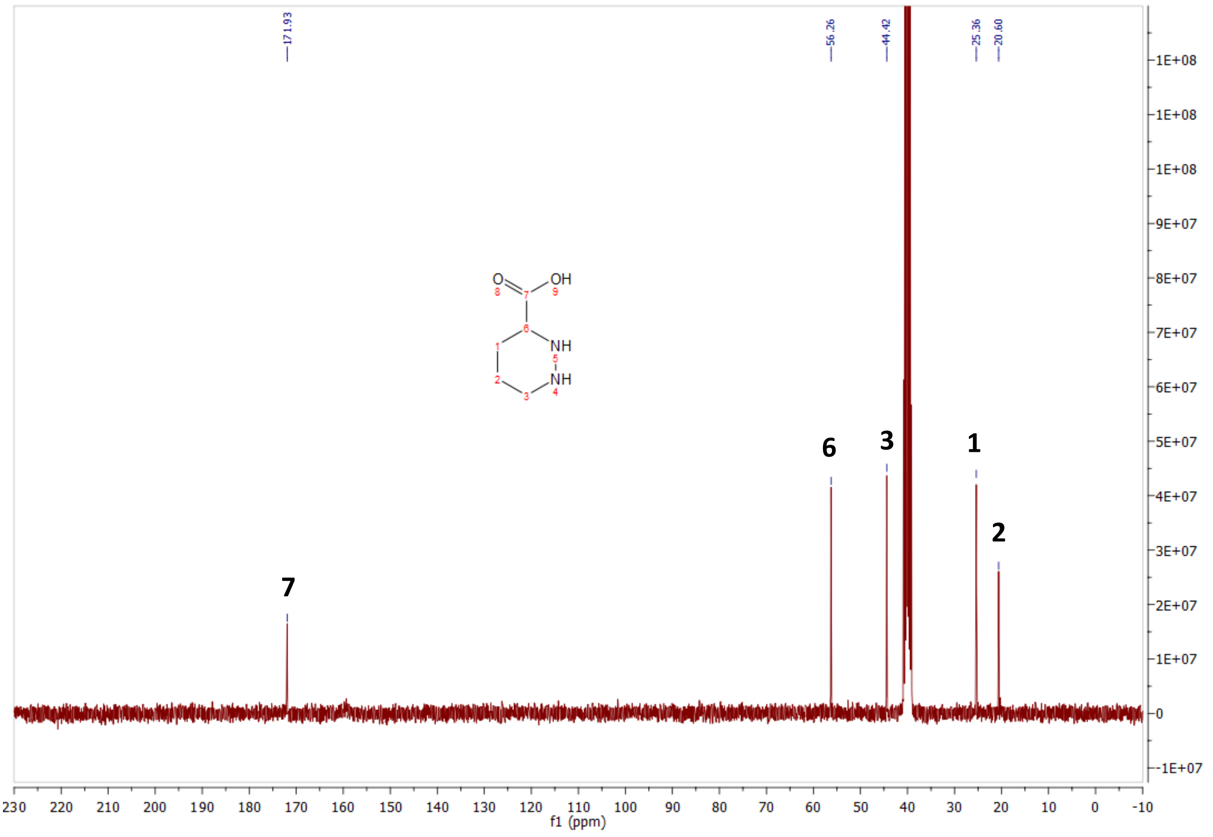

#### Boc-D-Nip-OSu (4) $^{1}$H NMR

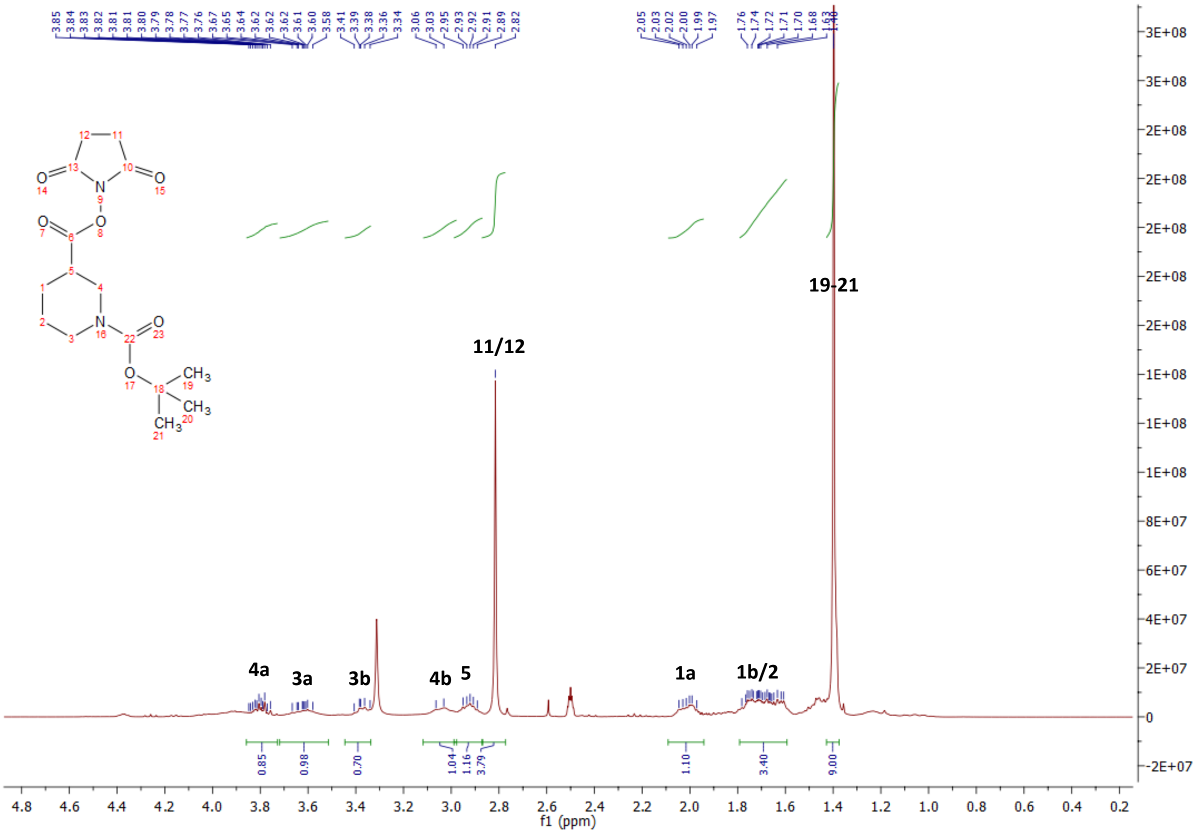

#### Boc-D-Nip-OSu (4) $^{13}$C NMR

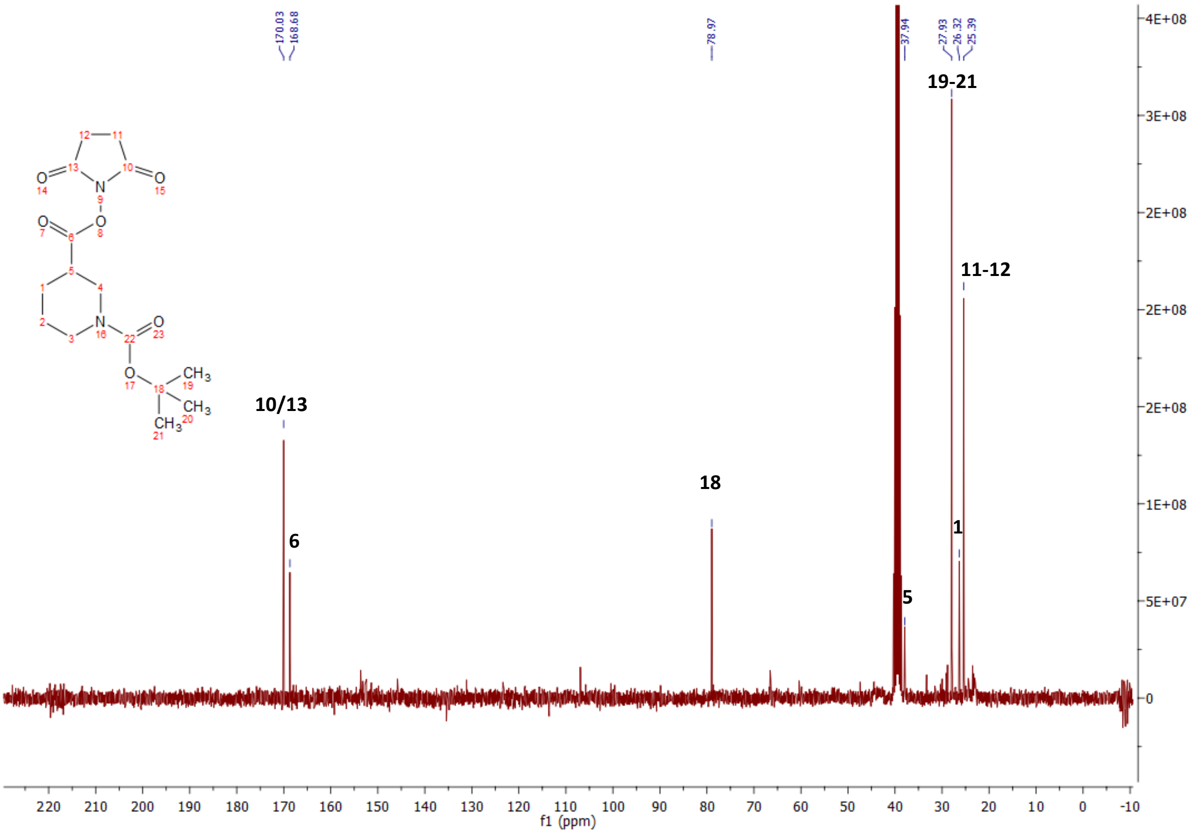

L-Pro-AMS (7) $^{1}$H NMR

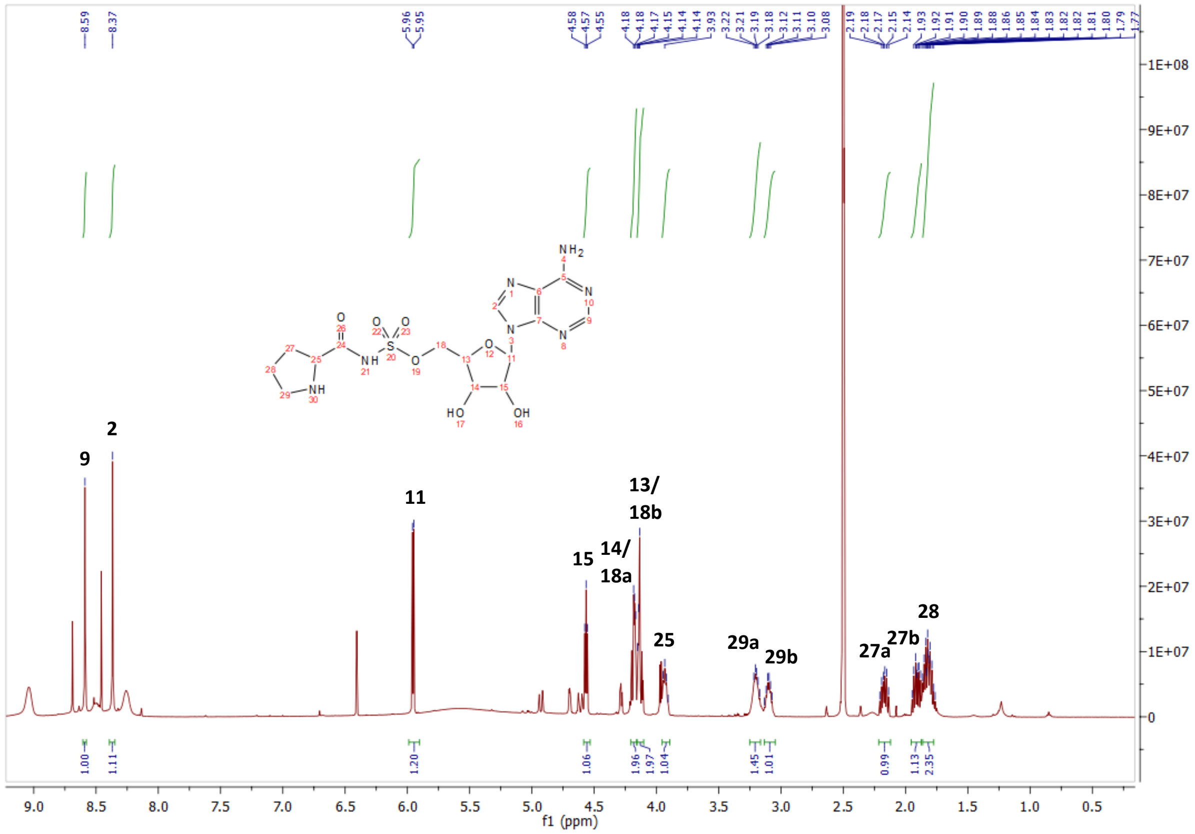

L-Pro-AMS (7) $^{13}$C NMR

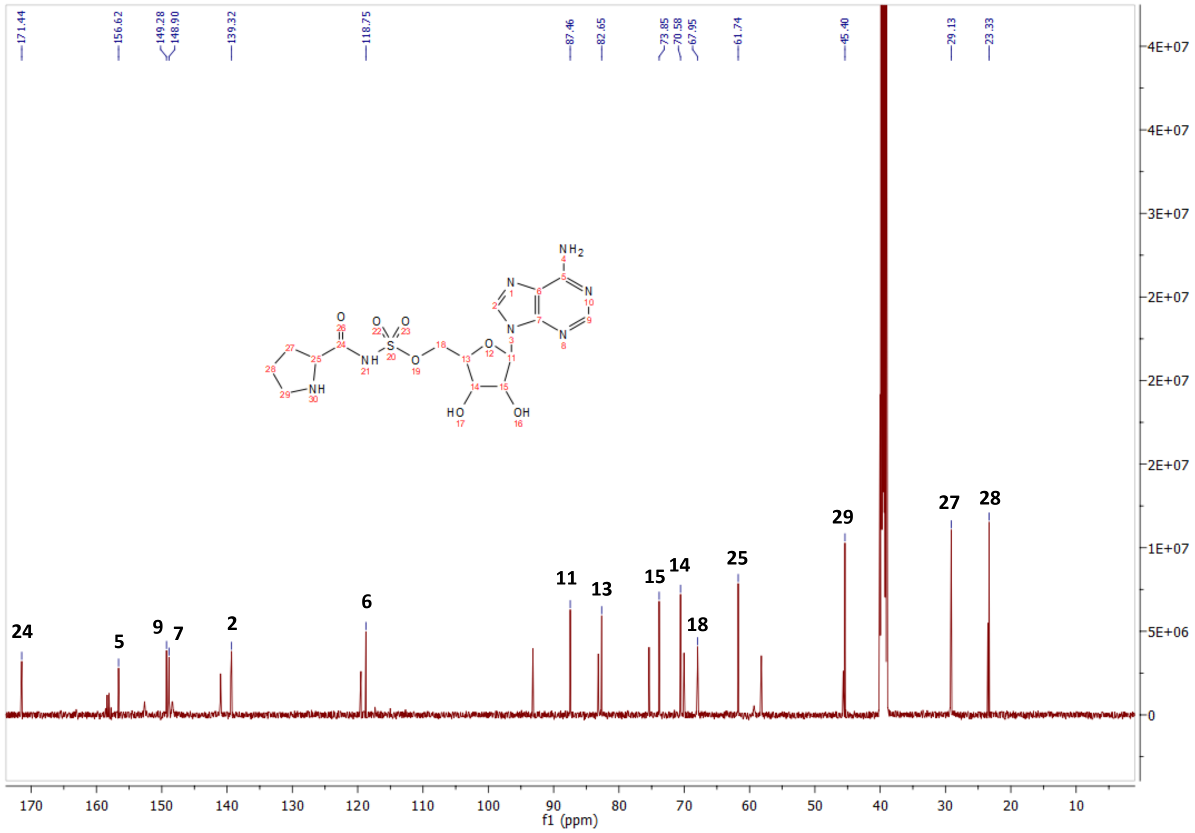

**L-Pro-AMS (7) COSY**

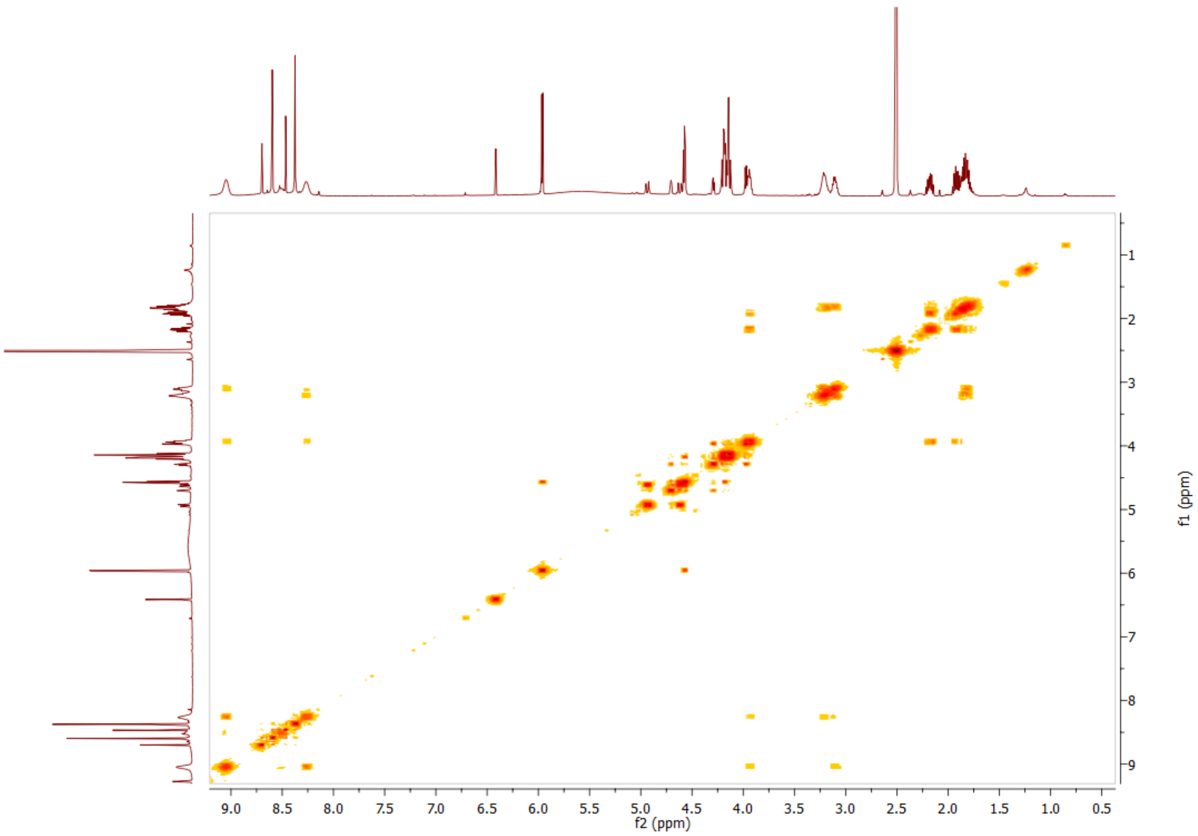

**L-Pro-AMS (7) HSQC**

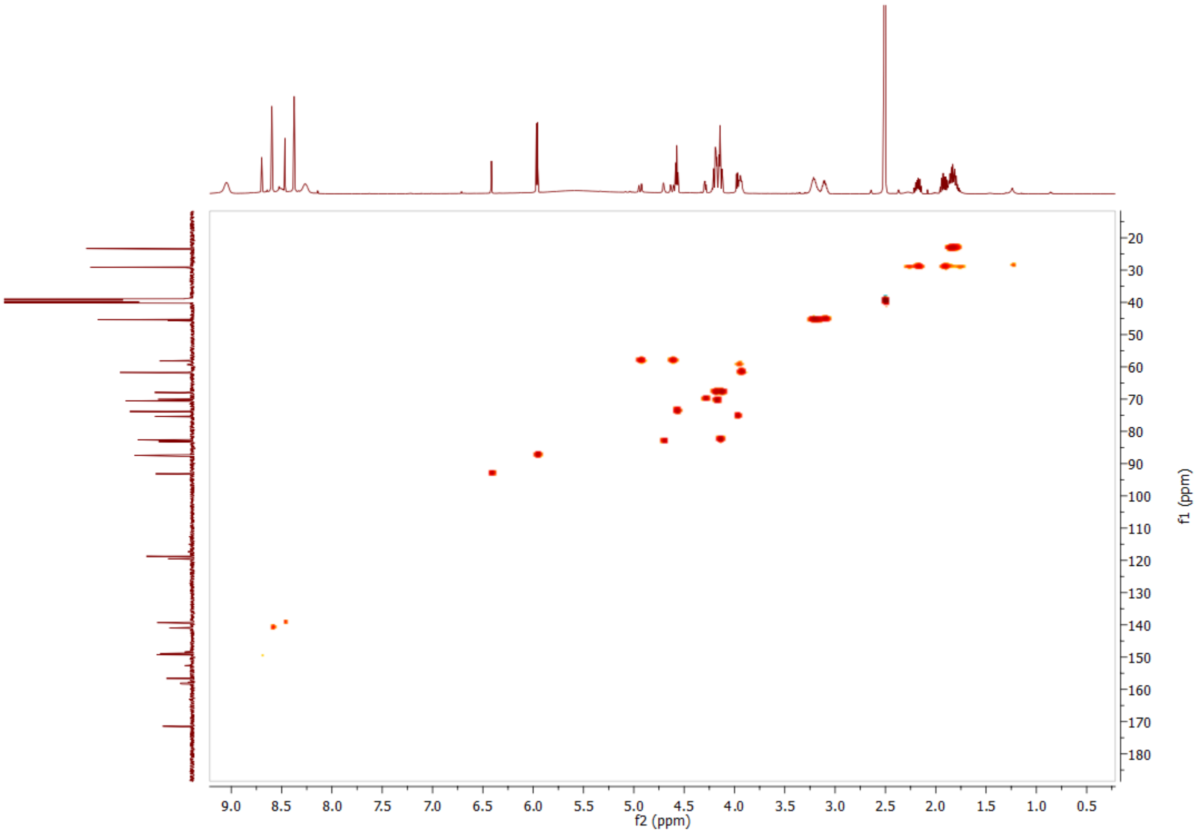

**D-Nip-AMS (8) $^{1}$H NMR**

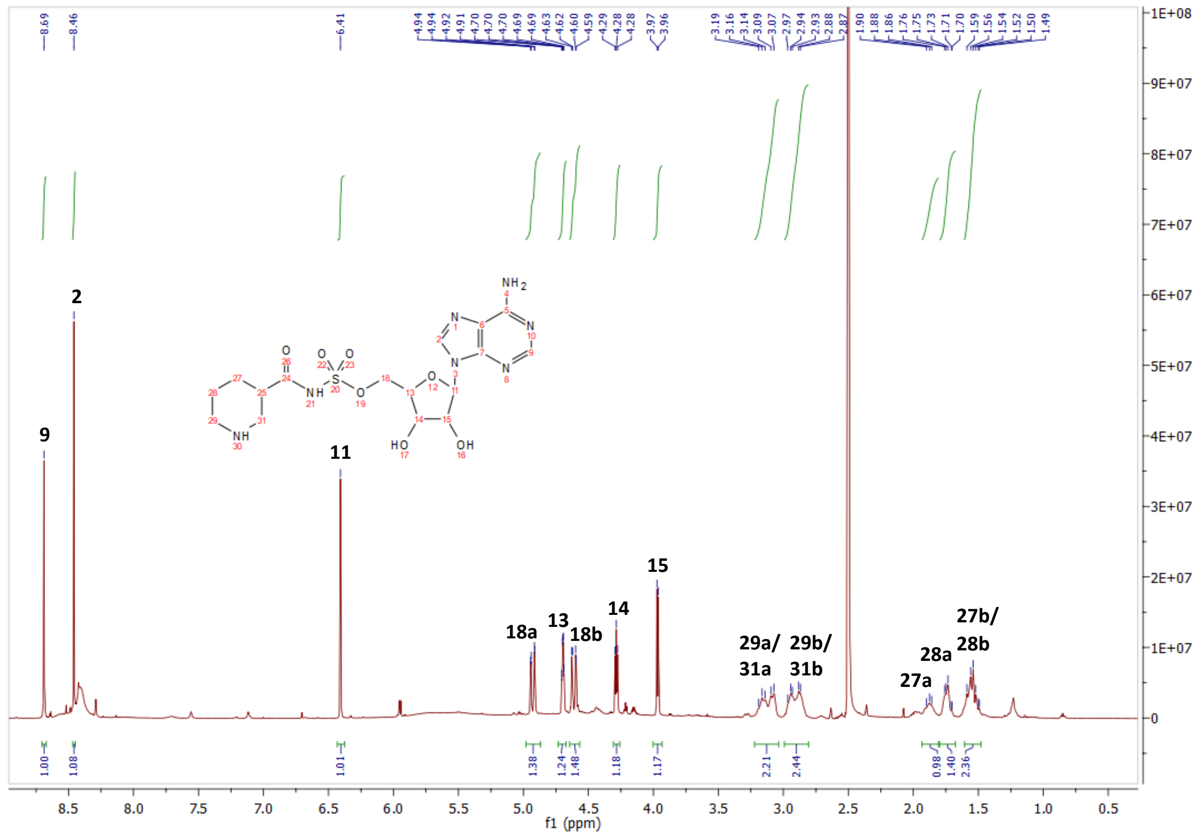

**D-Nip-AMS (8) $^{13}$C NMR**

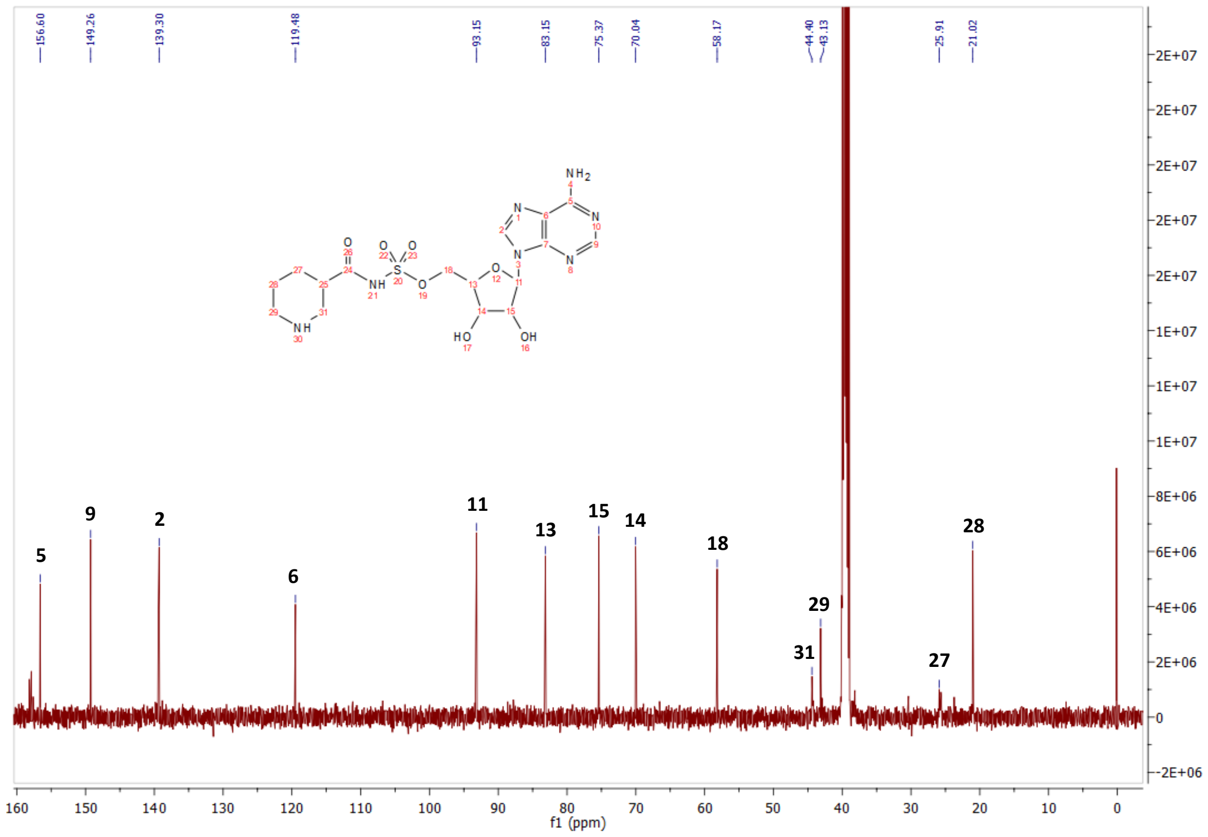

**D-Nip-AMS (8) COSY**

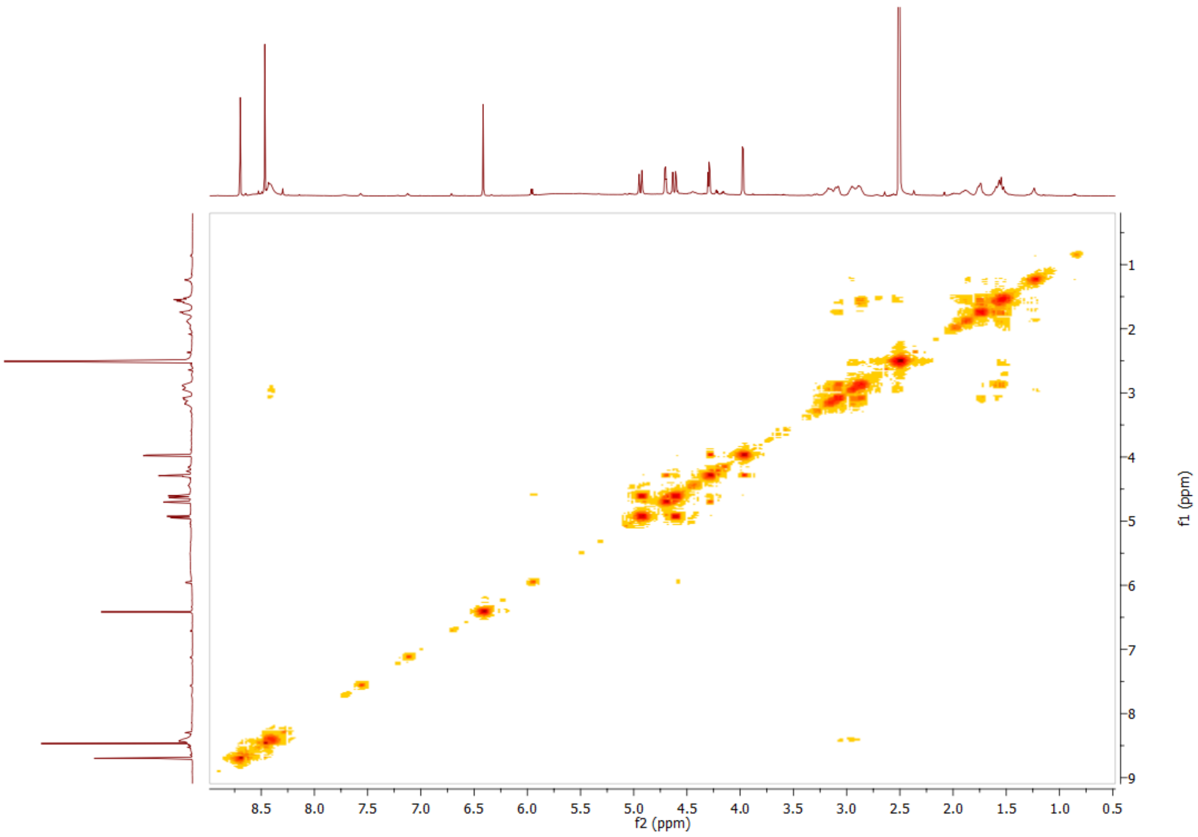

**D-Nip-AMS (8) HSQC**

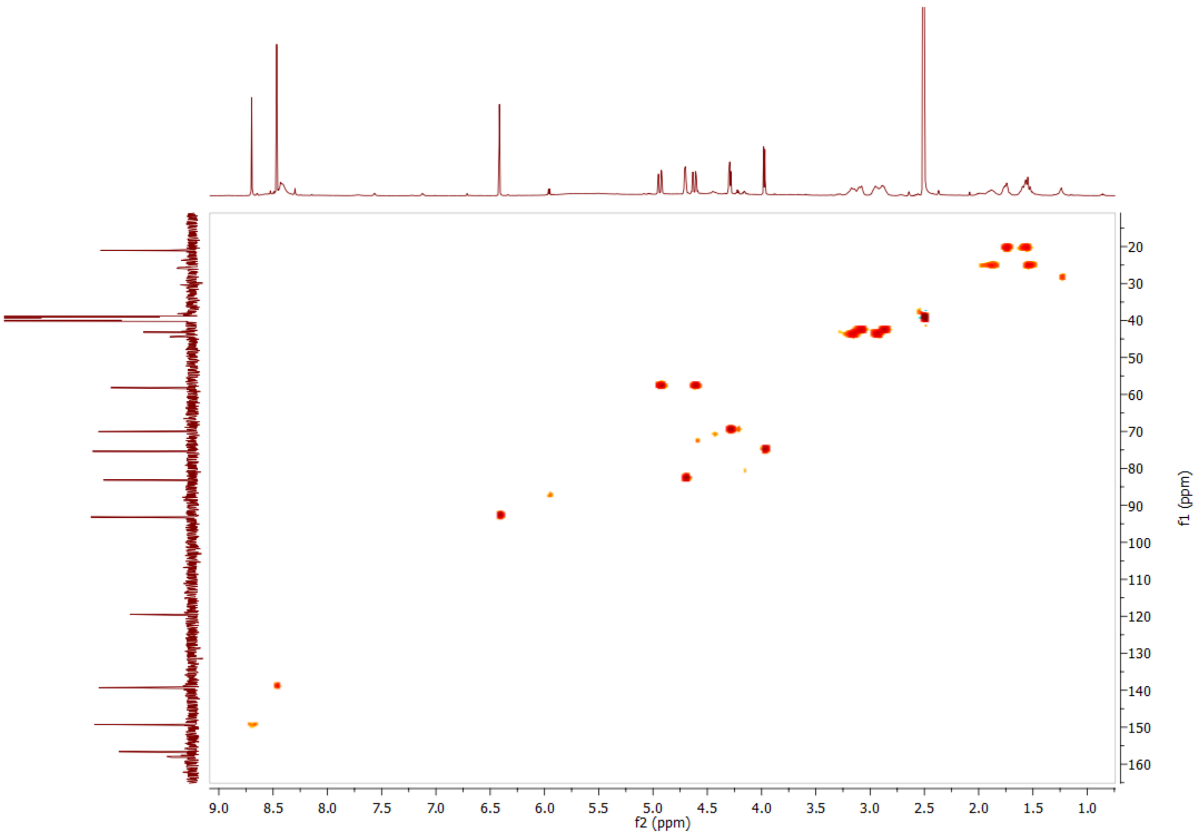

### 12. Protein sequences

Mutated positions are highlighted.

**GrsA**

MLNSSKSILIHAQNKNGTHEEEQYLFAVNNTKAEYPRDKTIHQLFEEQVSKRPNNVAIVCENEQLTYHELNVKANQLARIFIE
KGIGKDTLVGIMMEKSIDLFIGILAVLKAGGAYVPIDIEYPKERIQYILDDSQARMLLTQKHLVHLIHNIQFNGQVEIFEEDT
IKIREGTNLHVPSKSTDLAYVIYTSGTTGNPKGTMLEHKGISNLKVFFENSLNVTEKDRIGQFASISFDASVWEMFMALLTGA
SLYIILKDTINDFVKFEQYINQKEITVITLPPTYVVHLDPERILSIQTLITAGSATSPSLVNKWKEKVTYINAYGPTETTICA
TTWVATKETIGHSVPIGAPIQNTQIYIVDENLQLKSVGEAGELCIGGEGLARGYWKRPELTSQKFVDNPFVPGEKLYKTGDQA
RWLSDGNIEYLGRIDNQVKIRGHRVELEEVESILLKHMYISETAVSVHKDHQEQPYLCAYFVSEKHIPLEQLRQFSSEELPTY
MIPSYFIQLDKMPLTSNGKIDRKQLPEPDLTFGMRVDYEAPRNEIEETLVTIWQDVLGIEKIGIKDNFYALGGDSIKAIQVAA
RLHSYQLKLETKDLLKYPTIDQLVHYIKDSKRRSEQGIVEGEIGLTPIQHWFFEQQFTNMHHWNQSYMLYRPNGFDKEILLRV
FNKIVEHHDALRMIYKHHNGKIVQINRGLEGTLFDFYTFDLTANDNEQQVICEESARLQNSINLEVGPLVKIALFHTQNGDHL
FMAIHHLVVDGISWRILFEDLATAYEQAMHQQTIALPEKTDSFKDWSIELEKYANSELFLEEAEYWHHLNYYTENVQIKKDYV
TMNNKQKNIRYVGMELTIEETEKLLKNVNKAYRTEINDILLTALGFALKEWADIDKIVINLEGHGREEILEQMNIARTVGWFT
SQYPVVLDMQKSDDLSYQIKLMKENLRRIPNKGIGYEIFKYLTTEYLRPVLPFTLKPEINFNYLGQFDTDVKTELFTRSPYSM
GNSLGPDGKNNLSPEGESYFVLNINGFIEEGKLHITFSYNEQQYKEDTIQQLSRSYKQHLLAIIEHCVQKEDTELTPSDFSFK
ELELEEMDDIFDLLADSLTGSRSHHHHHH

**GrsB1**

MSTFKKEHVQDMYRLSPMQEGMLFHALLDKDKNAHLVQMSIAIEGIVDVELLSESLNILIDRYDVFRTTFLHEKIKQPLQVVL
KERPVQLQFKDISSLDEEKREQAIEQYKYQDGETVFDLTRDPLMRVAIFQTGKVNYQMIWSFHHILMDGWCFNIIFNDLFNIY
LSLKEKKPLQLEAVQPYKQFIKWLEKQDKQEALRYWKEHLMNYDQSVTLPKKKAAINNTTYEPAQFRFAFDKVLTQQLLRIAN
QSQVTLNIVFQTIWGIVLQKYNSTNDVVYGSVVSGRPSEISGIEKMVGLFINTLPLRIQTQKDQSFIELVKTVHQNVLFSQQH
EYFPLYEIQNHTELKQNLIDHIMVIENYPLVEELQKNSIMQKVGFTVRDVKMFEPTNYDMTVMVLPRDEISVRLDYNAAVYDI
DFIKKIEGHMKEVALCVANNPHVLVQDVPLLTKQEKQHLLVELHDSITEYPDKTIHQLFTEQVEKTPEHVAVVFEDEKVTYRE
LHERSNQLARFLREKGVKKESIIGIMMERSVEMIVGILGILKAGGAFVPIDPEYPKERIGYMLDSVRLVLTQRHLKDKFAFTK
ETIVIEDPSISHELTEEIDYINESEDLFYIIYTSGTTGKPKGVMLEHKNIVNLLHFTFEKTNINFSDKVLQYTTCSFDVCYQE
IFSTLLSGGQLYLIRKETQRDVEQLFDLVKRENIEVLSFPVAFLKFIFNEREFINRFPTCVKHIITAGEQLVVNNEFKRYLHE
HNVHLHNHYGPSETHVVTTYTINPEAEIPELPPIGKPISNTWIYILDQEQQLQPQGIVGELYISGANVGRGYLNNQELTAEKF
FADPFRPNERMYRTGDLARWLPDGNIEFLGRADHQVKIRGHRIELGEIEAQLLNCKGVKEAVVIDKADDKGGKYLCAYVVMEV
EVNDSELREYLGKALPDYMIPSFFVPLDQLPLTPNGKIDRKSLPNLEGIVNTNAKYVVPTNELEEKLAKIWEEVLGISQIGIQ
DNFFSLGGHSLKAITLISRMNKECNVDIPLRLLFEAPTIQEISNYINGGSRSHHHHHH

**GrsB1-AYA**

MSTFKKEHVQDMYRLSPMQEGMLFHALLDKDKNAHLVQMSIAIEGIVDVELLSESLNILIDRYDVFRTTFLHEKIKQPLQVVL
KERPVQLQFKDISSLDEEKREQAIEQYKYQDGETVFDLTRDPLMRVAIFQTGKVNYQMIWSFHHILMDGWCFNIIFNDLFNIY
LSLKEKKPLQLEAVQPYKQFIKWLEKQDKQEALRYWKEHLMNYDQSVTLPKKKAAINNTTYEPAQFRFAFDKVLTQQLLRIAN
QSQVTLNIVFQTIWGIVLQKYNSTNDVVYGSVVSGRPSEISGIEKMVGLFINTLPLRIQTQKDQSFIELVKTVHQNVLFSQQH
EYFPLYEIQNHTELKQNLIDHIMVIENYPLVEELQKNSIMQKVGFTVRDVKMFEPTNYDMTVMVLPRDEISVRLDYNAAVYDI
DFIKKIEGHMKEVALCVANNPHVLVQDVPLLTKQEKQHLLVELHDSITEYPDKTIHQLFTEQVEKTPEHVAVVFEDEKVTYRE
LHERSNQLARFLREKGVKKESIIGIMMERSVEMIVGILGILKAGGAFVPIDPEYPKERIGYMLDSVRLVLTQRHLKDKFAFTK
ETIVIEDPSISHELTEEIDYINESEDLFYIIYTSGTTGKPKGVMLEHKNIVNLLHFTFEKTNINFSDKVLQYTTCSFDVCYQE
IFSTLLSGGQLYLIRKETQRDVEQLFDLVKRENIEVLSFPVAFLKFIFNEREFINRFPTCVKHIITAGEQLVVNNEFKRYLHE
HNVHLHNAYGPSETHYATTYTINPEAEIPELPPIGKPISNTWIYILDQEQQLQPQGIVGELYISGANVGRGYLNNQELTAEKF
FADPFRPNERMYRTGDLARWLPDGNIEFLGRADHQVKIRGHRIELGEIEAQLLNCKGVKEAVVIDKADDKGGKYLCAYVVMEV
EVNDSELREYLGKALPDYMIPSFFVPLDQLPLTPNGKIDRKSLPNLEGIVNTNAKYVVPTNELEEKLAKIWEEVLGISQIGIQ
DNFFSLGGHSLKAITLISRMNKECNVDIPLRLLFEAPTIQEISNYINGGSRSHHHHHH

**GrsB1-AYA-SQSF**

MSTFKKEHVQDMYRLSPMQEGMLFHALLDKDKNAHLVQMSIAIEGIVDVELLSESLNILIDRYDVFRTTFLHEKIKQPLQVVL
KERPVQLQFKDISSLDEEKREQAIEQYKYQDGETVFDLTRDPLMRVAIFQTGKVNYQMIWSFHHILMDGWCFNIIFNDLFNIY
LSLKEKKPLQLEAVQPYKQFIKWLEKQDKQEALRYWKEHLMNYDQSVTLPKKKAAINNTTYEPAQFRFAFDKVLTQQLLRIAN
QSQVTLNIVFQTIWGIVLQKYNSTNDVVYGSVVSGRPSEISGIEKMVGLFINTLPLRIQTQKDQSFIELVKTVHQNVLFSQQH
EYFPLYEIQNHTELKQNLIDHIMVIENYPLVEELQKNSIMQKVGFTVRDVKMFEPTNYDMTVMVLPRDEISVRLDYNAAVYDI
DFIKKIEGHMKEVALCVANNPHVLVQDVPLLTKQEKQHLLVELHDSITEYPDKTIHQLFTEQVEKTPEHVAVVFEDEKVTYRE
LHERSNQLARFLREKGVKKESIIGIMMERSVEMIVGILGILKAGGAFVPIDPEYPKERIGYMLDSVRLVLTQRHLKDKFAFTK
ETIVIEDPSISHELTEEIDYINESEDLFYIIYTSGTTGKPKGVMLEHKNIVNLLHFTFEKTNINFSDKVLQYTTCSFDVCYQE
IFSTLLSGGQLYLIRKETQRDVEQLFDLVKRENIEVLSFPVAFLKFIFNEREFINRFPTCVKHIISAGEQLVVNNEFKRYLHE
HNVHLHNAYGQSESFYATTYTINPEAEIPELPPIGKPISNTWIYILDQEQQLQPQGIVGELYISGANVGRGYLNNQELTAEKF
FADPFRPNERMYRTGDLARWLPDGNIEFLGRADHQVKIRGHRIELGEIEAQLLNCKGVKEAVVIDKADDKGGKYLCAYVVMEV

EVNDSELREYLGKALPDYMIPSFFVPLDQLPLTPNGKIDRKSLPNLEGIVNTNAKYVVPTNELEEKLAKIWEEVLGISQIGIQ
DNFFSLGGHSLKAITLISRMNKECNVDIPLRLLFEAPTIQEISNYINGGSRSHHHHHH

**GrsB1-AYA-SQSF-VM**

MSTFKKEHVQDMYRLSPMQEGMLFHALLDKDKNAHLVQMSIAIEGIVDVELLSESLNILIDRYDVFRTTFLHEKIKQPLQVVL
KERPVQLQFKDISSLDEEKREQAIEQYKYQDGETVFDLTRDPLMRVAIFQTGKVNYQMIWSFHHILMDGWCFNIIFNDLFNIY
LSLKEKKPLQLEAVQPYKQFIKWLEKQDKQEALRYWKEHLMNYDQSVTLPKKKAAINNTTYEPAQFRFAFDKVLTQQLLRIAN
QSQVTLNIVFQTIWGIVLQKYNSTNDVVYGSVVSGRPSEISGIEKMVGLFINTLPLRIQTQKDQSFIELVKTVHQNVLFSQQH
EYFPLYEIQNHTELKQNLIDHIMVIENYPLVEELQKNSIMQKVGFTVRDVKMFEPTNYDMTVMVLPRDEISVRLDYNAAVYDI
DFIKKIEGHMKEVALCVANNPHVLVQDVPLLTKQEKQHLLVELHDSITEYPDKTIHQLFTEQVEKTPEHVAVVFEDEKVTYRE
LHERSNQLARFLREKGVKKESIIGIMMERSVEMIVGILGILKAGGAFVPIDPEYPKERIGYMLDSVRLVLTQRHLKDKFAFTK
ETIVIEDPSISHELTEEIDYINESEDLFYIIYTSGTTGKPKGVMLEHKNIVNVLHFTFEKTNINFSDKVLQYTTCSFDVCYQE
IFSTLLSGGQLYLIRKETQRDVEQLFDLVKRENIEVLSMPVAFLKFIFNEREFINRFPTCVKHIISAGEQLVVNNEFKRYLHE
HNVHLHNAYGQSESFYATTYTINPEAEIPELPPIGKPISNTWIYILDQEQQLQPQGIVGELYISGANVGRGYLNNQELTAEKF
FADPFRPNERMYRTGDLARWLPDGNIEFLGRADHQVKIRGHRIELGEIEAQLLNCKGVKEAVVIDKADDKGGKYLCAYVVMEV
EVNDSELREYLGKALPDYMIPSFFVPLDQLPLTPNGKIDRKSLPNLEGIVNTNAKYVVPTNELEEKLAKIWEEVLGISQIGIQ
DNFFSLGGHSLKAITLISRMNKECNVDIPLRLLFEAPTIQEISNYINGGSRSHHHHHH

**GrsB1-AYA-SQSF-VM-MWG**

MSTFKKEHVQDMYRLSPMQEGMLFHALLDKDKNAHLVQMSIAIEGIVDVELLSESLNILIDRYDVFRTTFLHEKIKQPLQVVL
KERPVQLQFKDISSLDEEKREQAIEQYKYQDGETVFDLTRDPLMRVAIFQTGKVNYQMIWSFHHILMDGWCFNIIFNDLFNIY
LSLKEKKPLQLEAVQPYKQFIKWLEKQDKQEALRYWKEHLMNYDQSVTLPKKKAAINNTTYEPAQFRFAFDKVLTQQLLRIAN
QSQVTLNIVFQTIWGIVLQKYNSTNDVVYGSVVSGRPSEISGIEKMVGLFINTLPLRIQTQKDQSFIELVKTVHQNVLFSQQH
EYFPLYEIQNHTELKQNLIDHIMVIENYPLVEELQKNSIMQKVGFTVRDVKMFEPTNYDMTVMVLPRDEISVRLDYNAAVYDI
DFIKKIEGHMKEVALCVANNPHVLVQDVPLLTKQEKQHLLVELHDSITEYPDKTIHQLFTEQVEKTPEHVAVVFEDEKVTYRE
LHERSNQLARFLREKGVKKESIIGIMMERSVEMIVGILGILKAGGAFVPIDPEYPKERIGYMLDSVRLVLTQRHLKDKFAFTK
ETIVIEDPSISHELTEEIDYINESEDLFYIIYTSGTTGKPKGVMLEHKNIVNVLHFTFEKTNINFSDKVLQYTTCSFDMCYWE
IFSTLLSGGQLYLIRKETQRDVEQLFDLVKRENIEVLSMPVAFLKFIFNEREFINRFPTCVKHIISGGEQLVVNNEFKRYLHE
HNVHLHNAYGQSESFYATTYTINPEAEIPELPPIGKPISNTWIYILDQEQQLQPQGIVGELYISGANVGRGYLNNQELTAEKF
FADPFRPNERMYRTGDLARWLPDGNIEFLGRADHQVKIRGHRIELGEIEAQLLNCKGVKEAVVIDKADDKGGKYLCAYVVMEV
EVNDSELREYLGKALPDYMIPSFFVPLDQLPLTPNGKIDRKSLPNLEGIVNTNAKYVVPTNELEEKLAKIWEEVLGISQIGIQ
DNFFSLGGHSLKAITLISRMNKECNVDIPLRLLFEAPTIQEISNYINGGSRSHHHHHH

**GrsB1-A$_{core}$**

MAHHHHHHSSGLEVLFQGPDSITEYPDKTIHQLFTEQVEKTPEHVAVVFEDEKVTYRELHERSNQLARFLREKGVKKESIIGI
MMERSVEMIVGILGILKAGGAFVPIDPEYPKERIGYMLDSVRLVLTQRHLKDKFAFTKETIVIEDPSISHELTEEIDYINESE
DLFYIIYTSGTTGKPKGVMLEHKNIVNLLHFTFEKTNINFSDKVLQYTTCSFDVCYQEIFSTLLSGGQLYLIRKETQRDVEQL
FDLVKRENIEVLSFPVAFLKFIFNEREFINRFPTCVKHIITAGEQLVVNNEFKRYLHEHNVHLHNHYGPSETHVVTTYTINPE
AEIPELPPIGKPISNTWIYILDQEQQLQPQGIVGELYISGANVGRGYLNNQELTAEKFFADPFRPNERMYRTGDLARWLPDGN
IEFLGRA

**MWG-A$_{core}$**

MAHHHHHHSSGLEVLFQGPDSITEYPDKTIHQLFTEQVEKTPEHVAVVFEDEKVTYRELHERSNQLARFLREKGVKKESIIGI
MMERSVEMIVGILGILKAGGAFVPIDPEYPKERIGYMLDSVRLVLTQRHLKDKFAFTKETIVIEDPSISHELTEEIDYINESE
DLFYIIYTSGTTGKPKGVMLEHKNIVNVLHFTFEKTNINFSDKVLQYTTCSFDMCYWEIFSTLLSGGQLYLIRKETQRDVEQL
FDLVKRENIEVLSMPVAFLKFIFNEREFINRFPTCVKHIISGGEQLVVNNEFKRYLHEHNVHLHNAYGQSESFYATTYTINPE
AEIPELPPIGKPISNTWIYILDQEQQLQPQGIVGELYISGANVGRGYLNNQELTAEKFFADPFRPNERMYRTGDLARWLPDGN
IEFLGRA

### 13. Supporting references

[1] S. Gruenewald, H. D. Mootz, P. Stehmeier, T. Stachelhaus, *Appl Environ Microbiol* **2004**, *70*, 3282–3291.

[2] A. Stanišić, A. Hüsken, P. Stephan, D. L. Niquille, J. Reinstein, H. Kries, *ACS Catal.* **2021**, *11*, 8692–8700.

[3] D. L. Niquille, D. A. Hansen, T. Mori, D. Fercher, H. Kries, D. Hilvert, *Nature Chem* **2018**, *10*, 282–287.

[4] F. Pourmasoumi, S. Hengoju, K. Beck, P. Stephan, L. Klopfleisch, M. Hoernke, M. A. Rosenbaum, H. Kries, *Biorxiv*, **2023**.

[5] T. Stachelhaus, D. Mootz, A. Marahiel, *Chemistry & Biology* **1999**, *6*, 493–505.

[6] G. L. Challis, J. Ravel, C. A. Townsend, *Chemistry & biology* **2000**, *7*, 211–224.

[7] K. Miyazaki, F. H. Arnold, *J Mol Evol* **1999**, *49*, 716–720.

[8] D. J. Wilson, C. C. Aldrich, *Analytical Biochemistry* **2010**, *404*, 56–63.

[9] W. Kabsch, *Acta Crystallogr D Biol Crystallogr* **2010**, *66*, 133–144.

[10] P. R. Evans, G. N. Murshudov, *Acta Crystallogr D Biol Crystallogr* **2013**, *69*, 1204–1214.

[11] D. Liebschner, P. V. Afonine, M. L. Baker, G. Bunkóczi, V. B. Chen, T. I. Croll, B. Hintze, L.-W. Hung, S. Jain, A. J. McCoy, N. W. Moriarty, R. D. Oeffner, B. K. Poon, M. G. Prisant, R. J. Read, J. S. Richardson, D. C. Richardson, M. D. Sammito, O. V. Sobolev, D. H. Stockwell, T. C. Terwilliger, A. G. Urzhumtsev, L. L. Videau, C. J. Williams, P. D. Adams, *Acta Crystallogr D Struct Biol* **2019**, *75*, 861–877.

[12] P. Emsley, K. Cowtan, *Acta Crystallogr D Biol Crystallogr* **2004**, *60*, 2126–2132.

[13] O. Trott, A. J. Olson, *J. Comput. Chem.* **2009**, NA-NA.

[14] J. Eberhardt, D. Santos-Martins, A. F. Tillack, S. Forli, *J. Chem. Inf. Model.* **2021**, *61*, 3891–3898.

[15] http://www.yasara.org/

[16] H. Land, M. S. Humble, in *Protein Engineering* (Eds.: U.T. Bornscheuer, M. Höhne), Springer New York, New York, NY, **2018**, pp. 43–67.

[17] E. F. Pettersen, T. D. Goddard, C. C. Huang, G. S. Couch, D. M. Greenblatt, E. C. Meng, T. E. Ferrin, *J. Comput. Chem.* **2004**, *25*, 1605–1612.

[18] The PyMOL Molecular Graphics System, Version 2.0 Schrödinger, LLC.

[19] E. Conti, T. Stachelhaus, M. a Marahiel, P. Brick, *Embo J* **1997**, *16*, 4174–4183.
